# Spectrotemporal signatures of driving and modulatory circuits across cortical and subcortical networks

**DOI:** 10.64898/2026.04.29.721627

**Authors:** M.N. O’Connell, A. Barczak, C. A. Mackey, T. McGinnis, K. Mackin, J.F. Smiley, C Bleiwas, P. Lakatos, C.E. Schroeder

**Author notes:** **Corresponding authors:** Monica Noelle O’Connell, Ph.D.

## Abstract

Sensory processing depends on interactions between neural circuits that convey and regulate information across cortical and subcortical networks. Classical frameworks distinguish driving inputs, which transmit sensory content via suprathreshold activation, from modulatory inputs, which alter neuronal excitability without directly eliciting spiking. However, physiological signatures of these circuit types that generalize widely across distributed brain regions remain unclear. Here, we functionally differentiate driving and modulatory circuits in the awake macaque brain by jointly quantifying suprathreshold multiunit activity (MUA) and oscillatory phase coherence (inter-trial coherence, ITC) across eight cortical and thalamic structures during auditory, visual, and motor sampling conditions. Preferred sensory stimuli elicited broadband ITC increases accompanied by robust MUA, yielding relatively uniform spectral distributions across adjacent frequency bands, consistent with driving inputs. In contrast, non-preferred sensory and motor-related events produced narrowband, frequency-specific ITC modulation without concurrent firing, and was characterized by dominant peaks at stimulation or event rates, which is consistent with modulatory inputs. This narrowband ITC modulation is indicative of coordinated phase alignment, capable of dynamically regulating information transfer, mediated by driving inputs, across thalamocortical circuits. These response types were observed within individual regions, revealing two separable modes of neural activity. These findings identify distinct spectrotemporal signatures of driving and modulatory activity and demonstrate that subthreshold oscillatory modulation is a widespread mechanism for coordinating multisensory and motor influences on perception.

## INTRODUCTION

Sensory processing relies on the coordinated activity of distributed neural circuits that convey and transform information from the periphery to cortex. Within this hierarchy, connections between regions can differ profoundly in their effect on target neurons. According to the influential framework first proposed by Sherman and Guillery (1998) and Jones (1998a,1998b), “driving” inputs transmit the main informative content of a signal, generating suprathreshold activation and shaping receptive field properties, whereas “modulatory” inputs primarily adjust the responsiveness of target neurons without directly driving spiking (Jones, 2001; Lakatos et al., 2007; Schroeder and Lakatos, 2009). These circuit types are not restricted to a single hierarchical level; both cortical and thalamic structures can receive mixtures of driving and modulatory inputs that together determine how sensory events are encoded and integrated (Barczak, O’Connell, & Schroeder, 2023).

The distinction between driving and modulatory circuits is measurable *in vivo* due to their specific physiological signatures. Driving inputs are typically associated with large, temporally precise synaptic currents that produce clear evoked responses in both local field potentials (LFPs) and multiunit activity (MUA). In contrast, modulatory inputs often produce subthreshold changes in membrane potential through the phase resetting of ongoing neural oscillations (which reflect fluctuations of excitability in neuronal ensembles) (P Lakatos, Chen, O’Connell, Mills, & Schroeder, 2007; Peter Lakatos et al., 2009, 2020; O’Connell et al., 2020; Shah et al., 2004). Thus, modulatory inputs regulate firing probabilities rather than causing obvious changes in the firing rates of neuron ensembles. Such subthreshold effects are particularly evident in studies utilizing inter-trial coherence (ITC), which indexes the phase alignment of neural oscillations across repeated events. Whereas broadband ITC increases often accompany stimulus-evoked firing, narrowband or low-frequency ITC modulations can reflect subtle entrainment of neuronal excitability - signatures of modulatory influence (Barczak et al., 2019; Obleser & Kayser, 2019; Charles E Schroeder & Lakatos, 2009).

Oscillatory modulation plays a critical role in shaping how the brain synchronizes sensory processing across space and time. Rhythmic inputs from auditory or visual streams, for example, can entrain low-frequency oscillations in cortical areas, aligning the phase of neuronal excitability with expected stimulus timing (Besle et al., 2011; Haegens & Zion Golumbic, 2018; Peter Lakatos, Karmos, Mehta, Ulbert, & Schroeder, 2008; Charles E Schroeder & Lakatos, 2009). Through this mechanism, modulatory inputs can enhance or decrease the effectiveness of driving signals if they arrive at the optimal or suboptimal oscillatory phase, respectively. Therefore, the functional distinction between driving and modulatory circuits is complemented by a dynamic interaction: modulatory entrainment establishes temporal windows of high or low excitability, while driving inputs convey detailed sensory content within those windows.

Accumulating evidence indicates that sensory pathways are inherently multisensory (Cappe, Rouiller, & Barone, 2009; Ghazanfar & Schroeder, 2006) and are further influenced by internally generated motor signals. Inputs from non-preferred modalities or motor systems have been found to act through modulatory pathways, shaping oscillatory phase and gain rather than directly driving activity. For instance, visual and auditory cortices can exhibit phase resetting in response to stimuli outside their dominant modality (Kayser, Petkov, & Logothetis, 2008; P Lakatos et al., 2007; Peter Lakatos et al., 2009; Mégevand et al., 2020; Naue et al., 2011; Plass et al., 2019; Romei, Gross, & Thut, 2012), and both can be entrained by saccadic eye movements that predictively modulate sensory responsiveness (Barczak et al., 2019; Ito, Maldonado, & Grün, 2013; Leszczynski et al., 2023; O’Connell et al., 2020). These findings suggest that motor and cross-modal signals operate primarily via modulatory circuits that influence the timing and excitability of sensory networks rather than transmitting content-rich driving input (with some exceptions: (Brosch, Selezneva, & Scheich, 2005; Mackey, O’Connell, Hackett, Schroeder, & Kajikawa, 2024)).

The concept of driving versus modulatory circuits is reasonably posited to be a general property of the brain, though its physiological expression across a wide variety of cortical and subcortical areas remains incompletely characterized. Many prior studies have focused on single structures, which makes it difficult to assess how driving and modulatory mechanisms operate in parallel across distributed nodes within sensory hierarchies. In the thalamus, for example, primary relay nuclei such as the medial and lateral geniculate bodies (MGB, LGN) receive well-characterized driving inputs from peripheral sensory organs but also receive modulatory inputs from other subcortical structures, plus feedback from cortex (Barczak et al., 2023; (Sherman & Guillery, 2002, 2011). Similarly, cortical sensory areas integrate both bottom-up driving and modulatory inputs (Jones, 1998, 2001), and top-down modulatory influences that shape oscillatory dynamics (Gilbert & Li, 2013; Helfrich & Knight, 2016). Determining if and how these functional distinctions - driving versus modulatory - are reflected in the temporal and spectral characteristics of neuronal responses across regions is critical for understanding how the brain coordinates perception across modalities and hierarchical levels.

Here, we sought to identify physiological signatures of driving and modulatory circuits across auditory, visual, motor and multisensory structures in the awake macaque brain. By simultaneously recording local field potentials and multiunit activity in multiple cortical and thalamic regions, we compared neural responses within these regions to sensory stimuli of preferred and non-preferred modalities, and internally generated saccadic events. We reasoned that driving inputs would elicit strong suprathreshold activation, which would be expressed as a broadband response in the field potential, accompanied by MUA, and that modulatory inputs would primarily induce narrowband oscillatory phase alignment without significant population spiking. By examining these response patterns across multiple structures, we aimed to characterize how driving and modulatory signals are distributed within and across thalamocortical networks. We further hypothesized that modulatory input regulates the phase relationships of oscillatory activity across regions, thereby dynamically facilitating or impeding the transfer of sensory information through driving pathways. In this view, modulatory circuits do not merely synchronize local excitability but align, or misalign, the phases of distributed networks to control when and where driving inputs are most effective. By jointly quantifying suprathreshold and subthreshold responses across auditory and visual thalamic nuclei and cortical areas, our study provides new insights into the functional organization of driving and modulatory circuits in the awake primate brain. Understanding this balance is critical for explaining how neural systems integrate multisensory inputs, including auditory, visual, and motor-related signals, generate predictions about upcoming input, and support coherent perception in dynamic environments.

## MATERIALS AND METHODS

### Subjects

In the present study, eye position and electrophysiological data were recorded from 1 male and 5 female macaques (*Macaca mulatta*) weighing 4.5-10kg, who had been prepared surgically for chronic awake electrophysiological recordings. There were 207 recording sites distributed across 137 experimental sessions (38, 46, 13, 40, 14 and 56 sites from 19, 32, 9, 30, 10 and 37 sessions with macaques B, G, M, R, T and U respectively). Prior to surgery, each animal was adapted to a custom fitted primate chair and to the recording chamber. All procedures were approved in advance by the Institutional Animal Care and Use Committee of the Nathan Kline Institute.

### Surgery

Preparation of subjects for chronic awake intracortical recording was performed using aseptic techniques, under general anesthesia, as described previously (C E Schroeder et al., 1998). The tissue overlying the calvarium was resected and appropriate portions of the cranium were removed. The neocortex and overlying dura were left intact. To provide access to the brain and to promote an orderly pattern of sampling across the surface of the auditory areas, Polyetheretherketone (PEEK) recording chambers (Rogue Research Inc.) were positioned normal to the cortical surface of the superior temporal plane for orthogonal (i.e., perpendicular to the layers) penetration of auditory cortex and thalamic structures beneath, as determined by pre-surgical structural MRI. Together with a PEEK headpost (to permit painless head restraint), they were secured to the skull with ceramic screws and embedded in dental acrylic. We used MRI compatible materials to allow for post implant imaging. A recovery time of six weeks was allowed before we resumed behavioral training and began data collection.

### Electrophysiology

During the experiments, animals sat in a primate chair in a dim, isolated, electrically shielded, sound-attenuated chamber with head fixed in position, and were monitored with infrared cameras. Neuroelectric activity was obtained using linear array multi-contact electrodes (23 contacts, 100 µm intercontact spacing, Plexon Inc.) from 66 A1 (20, 10, 6, 16, 7 and 7 sites in macaques B, G, M, R, T and U respectively), 31 secondary auditory cortex (Belt), (6, 11, 2, 4, 2 and 6 sites in macaques B, G, M, R, T and U respectively), 16 ventral division of the medial geniculate body (2, 5, 1 and 8 sites in macaques G, R, T and U respectively), 34 nonlemniscal medial geniculate body (4, 9, 1, 8, 3 and 9 sites in macaques B, G, M, R, T and U respectively), 11 lateral geniculate nucleus (1, 2, 4 and 4 sites in macaques B, G, R and U respectively), 21 medial pulvinar (8, 1,1, 1 and 10 sites in macaques G, M, R, T and U respectively), 14 pulvinar (2, 4, 2, 1 and 5 sites in macaques B, G, M, R, and U respectively) and 14 motor cortex (5, 1, 1 and 7 sites in macaques B, M, R and U respectively) sites. The multielectrodes were inserted acutely through guide tube grid inserts, lowered through the dura into the brain, and positioned such that the electrode channels would span as much of a sub-cortical nucleus as 2.2mm would allow (see section below titled “Identification of Recording Structures” and Figure 1), and all layers of a cortical region. Final electrode position was guided by creating and inspecting current source density (CSD) and multiunit (MUA) response profiles to binaural broadband noise bursts and strobe flashes. Neuroelectric signals were impedance matched with a pre-amplifier (10x gain, bandpass dc-10 kHz) situated on the electrode, and after further amplification (500x) they were recorded continuously with a 0.01 - 8000 Hz bandpass digitizer with a sampling rate of 44 kHz and precision of 12-bits using the Alpha Omega SnR system. The signal was split into the field potential (0.1-300Hz) and MUA (300-5000Hz) range by zero phase shift digital filtering. MUA data was also rectified in order to improve the estimation of firing of the local neuronal ensemble (Legatt, Arezzo, & Vaughan, 1980). One-dimensional current source density (CSD) profiles were calculated from the local field potential profiles using a three-point formula for the calculation of the second spatial derivative of voltage (Freeman & Nicholson, 1975). The advantage of CSD profiles is that they are not affected by volume conduction like the local field potentials (Kajikawa & Schroeder, 2011, 2015), and they also provide a more direct index of the location, direction, and density of the net transmembrane current flow (Mitzdorf, 1985;Schroeder et al., 1998). Bipolar signals were derived by computing the difference between adjacent contacts along each linear electrode array. This spatial differentiation reduces the influence of volume-conducted activity and enhances sensitivity to locally generated signals. Bipolar derivations were used for analyses of subcortical recordings, where laminar current source density estimates are not applicable.

**Figure 1.**
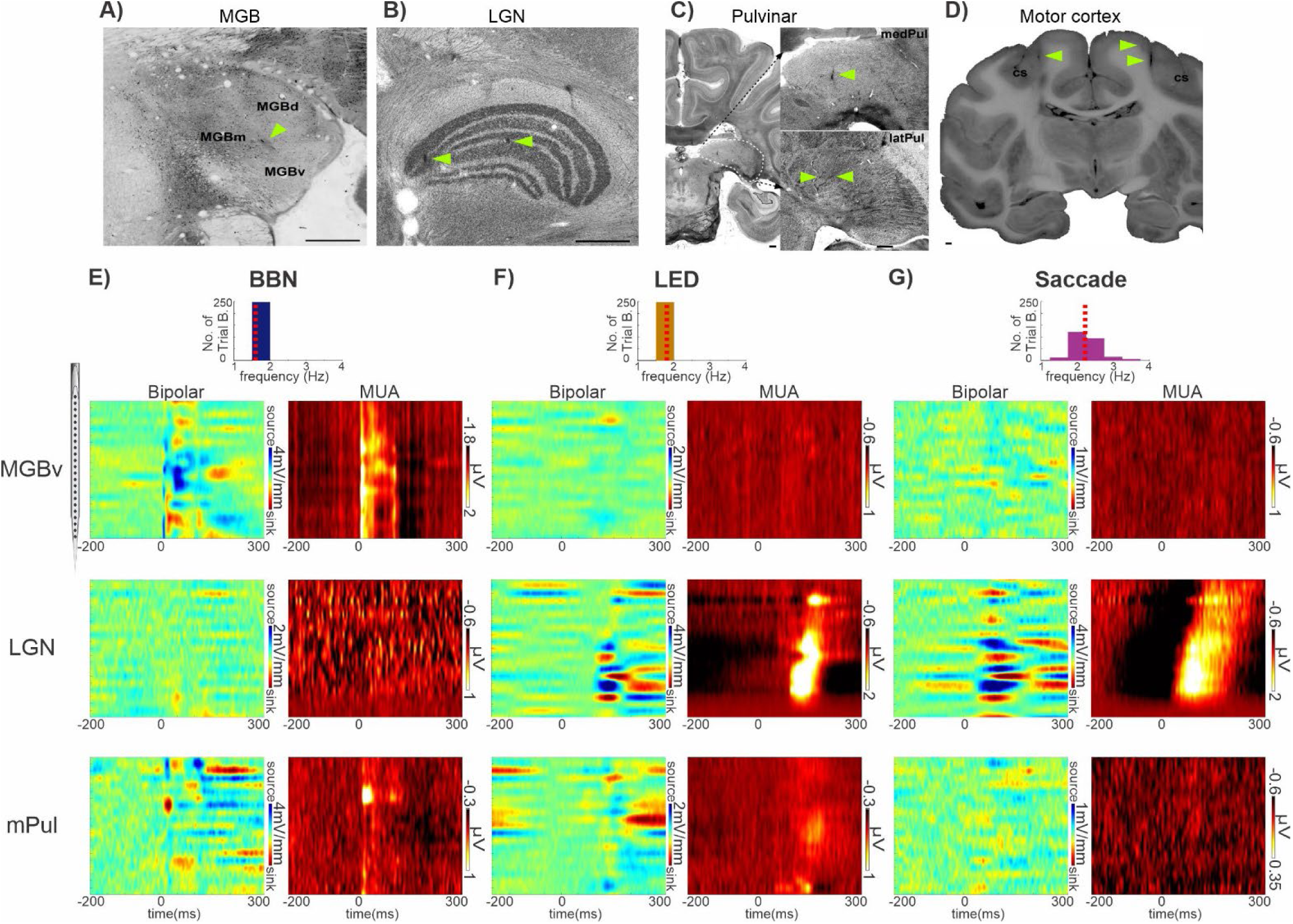
Representative MUA and bipolar field potential profiles associated with preferred and non-preferred modalities in sub-cortical structures. **A-D**) Histological verification of recording sites. (**A**) Calbindin staining of MGB (dorsal, medial and ventral subdivisions). (**B**) Nissl-stained section of LGN. (**C**) Nissl-stained sections of pulvinar, with higher magnification insets showing electrodes in medial and lateral subnuclei. (**D**) Block-face image taken during brain sectioning, showing electrodes in motor cortex just medial to the central sulcus (cs). Yellow arrows indicate electrode tracks. Panels A-C are oriented with the medial brain to the left. Scale bars = 1 mm. **E-G**) Representative bipolar field potential and MUA profiles from MGBv (top), LGN (middle), and mPul (bottom), with corresponding event-rate distributions (top panels; red dotted lines = median rates). **E)** BBN (1.6 Hz). **F)** LED (1.8 Hz; same sites). **G)** Saccades (2.2 Hz), aligned to onset. Note different amplitude scales.

### Eye Tracking and Saccade Detection

While the animal’s head was immobilized, eye position was monitored at a sampling rate of 500Hz using the Eyelink 1000 Plus system (SR Research Ltd). Vertical and horizontal eye position was transformed into changes in voltage and simultaneously recorded with the electrophysiological data. Using custom-written functions in Matlab (Mathworks, Natick, MA) we detected saccades if the magnitude of the eye movement (calculated as the length of the eye movement vector) exceeded an arbitrary cut-off value (not measured in distance since our eye tracking system was not calibrated). The threshold was set relatively high since we did not want to capture any artifacts (e.g. related to movement). Therefore, micro-saccades were not detected. The cut-off was optimized based on input from several lab members who have experience with manual saccade detection. Saccades had to last for at least 8ms and the minimum timing between saccades (i.e. minimum duration of fixation) had to be at least 120ms. Only resting state trial blocks with at a minimum of 40 verified saccades were included for analyses.

### Stimulus Presentation

At the beginning of each experimental session, streams of 50 dB SPL broadband noise (BBN) bursts and strobe flashes (Grass Instrument photo stimulator) were presented separately to elicit auditory and/or visual neural responses to locate brain structures and determine final electrode position. Once electrode position was refined, the frequency tuning of each recording site was determined using a “suprathreshold” method (Fu et al., 2004; C E Schroeder et al., 2001; Steinschneider, Reser, Schroeder, & Arezzo, 1995). The method entailed the presentation of a pseudorandom train of 14 different frequency pure tones ranging from 353.5Hz to 32kHz in half octave steps and a BBN burst at 50 dB SPL (duration: 100ms, rising/falling time: 4ms, stimulus onset asynchrony (SOA) = 319, n = 1600). Next, for the collection of the data used for the main analyses in this paper, subjects sat passively in a dim sound attenuated chamber through three conditions. Condition 1 included the the presentation of a rhythmic stream of BBN bursts at 50 dB SPL (duration: 100ms, r/f time: 4ms, SOA = 624ms, corresponding to a 1.6Hz repetition rate). Every BBN stream trial block consisted of approximately 100 BBN bursts and thus lasted just over 1 minute. Auditory stimuli were produced using Tucker Davis Technology’s System III coupled with two MF-1 free field speakers which were positioned 12 inches diagonally in front of each subject at ear level. Condition 2 included the presentation of a rhythmic stream of LED flashes which were 25ms in duration and had an SOA of 562ms (corresponding to a 1.8Hz repetition rate). Each LED stream trial block consisted of approximately 500 LED flashes and thus lasted almost 5 minutes. The LED was positioned 39 inches directly in front of the subject at eye level. In a subset of experiments (n= 33), the LED stream was presented at a theta rate of 6Hz (SOA of 166ms). Condition 3 included the absence of any stimuli (resting state condition) during which saccades were detected. The resting state trial block lasted approximately 5 minutes. There was no behavioral task in any condition and thus no behaviour was required of the NHP. Animals were kept alert by interacting with them prior to and after each trial block.

### Identification of Recording Structures

Consultation of the Paxinos macaque anatomical atlas with stereotaxic co-ordinates guided the location of each electrode penetration, along with structural MRI of each macaque. Recording locations were additionally verified histologically, as described below. Auditory cortical recording sites were initially identified based on location in the recording chamber, depth from superior surface of the brain and responsiveness to auditory stimuli. Primary (A1) and secondary (Belt) auditory cortical recording sites were separated based on the sharpness of frequency tuning, the inspection of the tonotopic progression across adjacent sites, onset latencies and relative sensitivity to pure tones versus broad-band noise of equivalent intensity (Peter Lakatos, Pincze, et al., 2005; Merzenich & Brugge, 1973; Rauschecker, Tian, Pons, & Mishkin, 1997). Motor cortical recording sites were characterized based on their more anterior position in the recording chamber, their shallow depth from the top of dura, the inversion of local field potentials to auditory stimuli and lack of frequency tuning.

Thalamic and subcortical structures were, at first, determined based on depth from superior surface of the brain and primary responsiveness to either auditory or visual stimuli (e.g. MGB and lateral geniculate nucleus (LGN)). The classification of recording sites located in the ventral division of MGB (MGBv) as opposed to in the dorsal and ventral subdivisions of the nucleus (MGB) was determined by comparing MUA response onset latencies to BBN and pure tone stimuli, and sharpness of frequency tuning curves. The ability to separately identify lateral pulvinar and LGN recording sites was based on recording depth, onset latencies of MUA responses to visual stimuli and medial-lateral location relative to other identified structures in the recording chamber. Medial pulvinar recording sites were labelled as such due to their recording depth and responsiveness to both auditory and visual stimuli. At the end of each animal’s experimental participation, functional assignment of the recording sites was confirmed histologically (Fig 1A-D) (Lakatos et al. 2020; (Barczak et al., 2018); Schroeder et al., 2001).

### Data analysis

Data were analyzed offline using native and custom-written functions in Matlab (Mathworks, Natick, MA). To identify suprathreshold event-related MUA responses at each electrode contact, we statistically compared the post-event (0-300ms) activity of “real” MUA epochs - time-locked to the event of interest (e.g., BBN onset) - with that of “control” MUA epochs using a Wilcoxon rank-sum test. Bonferroni correction was applied to account for multiple comparisons across both recording contacts within each structure and timepoints within each analysis window. An evoked response was defined as a period of at least 4 consecutive milliseconds in which MUA activity significantly differed between real and control epochs. Control MUA epochs were generated by randomly selecting timepoints within each trial block to serve as pseudo-event onsets, with the number of random timepoints matched to the number of real event onsets in that block. For each structure, if multiple linear-array contacts exhibited significant evoked MUA responses, the contact whose significant response was closest to event onset (0ms) - whether excitatory or inhibitory - was selected for display in Figure 2.

**Figure 2.**
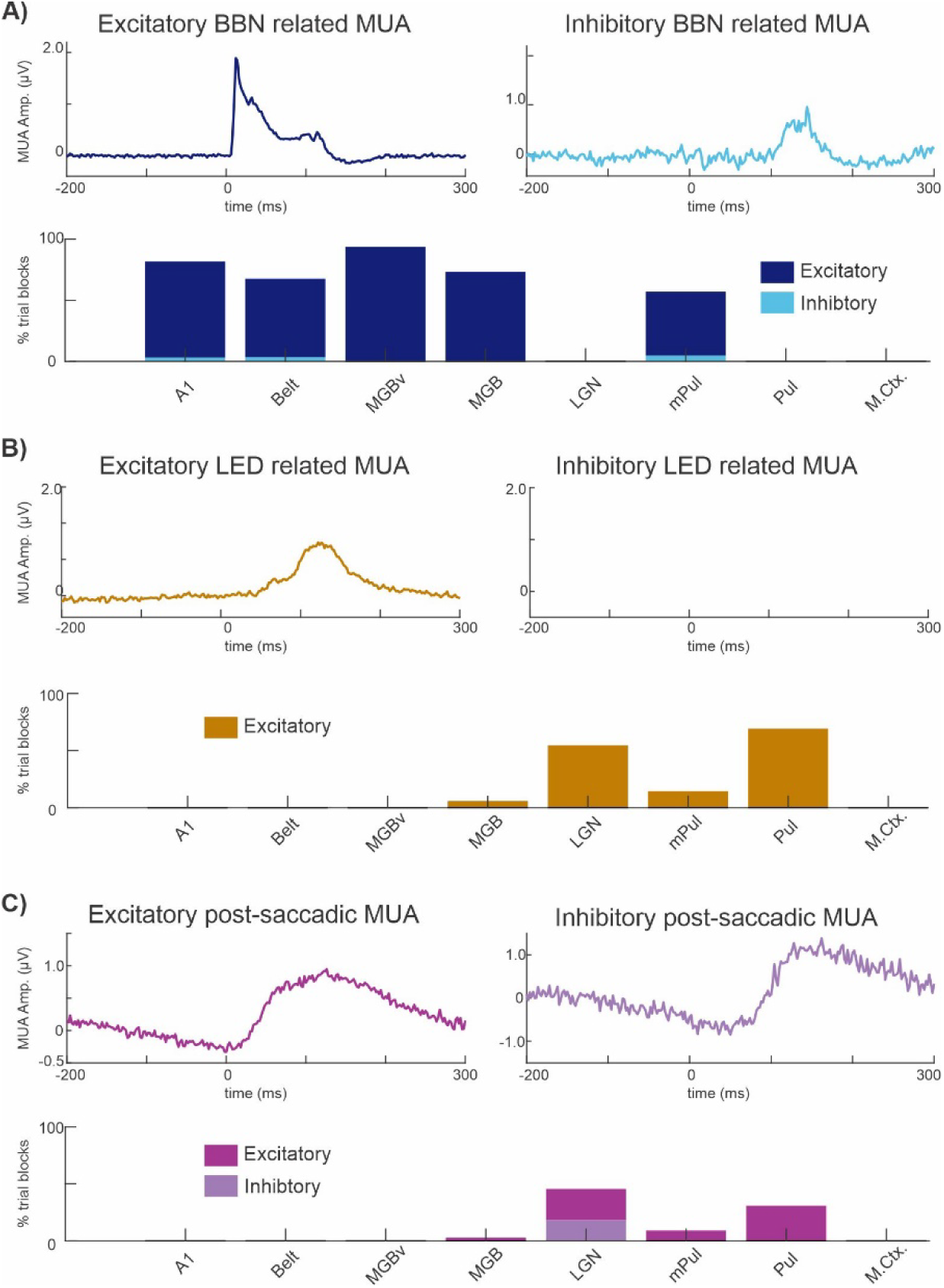
Excitatory and inhibitory suprathreshold MUA responses across 8 cortical and subcortical brain structures. **A-C)** Average MUA waveforms and corresponding bar graphs summarizing the proportion of recording sites exhibiting significant evoked responses (0–300 ms) across brain regions. Left and right traces show excitatory and inhibitory responses, respectively, color-coded by condition. Bar graphs indicate the percentage of sites within each region exhibiting each response type. **A)** BBN (navy: excitatory; cyan: inhibitory). **B)** LED (mustard: excitatory; no inhibitory responses). **C)** Saccades (purple: excitatory; light purple: inhibitory).

To quantify the phase coherence of neuronal oscillatory activity across trials, the instantaneous phase of each single trial signal was extracted from MUA, CSD signals for cortical sites and from bipolar derivations for subcortical sites using Morlet wavelet decomposition across 69 frequency scales ranging from 0.7 to 50 Hz. ITC values range from 0 to 1; higher values indicate that the observations (oscillatory phase at a given time-point across trials) are clustered more closely around the mean than lower values (phase distribution is random). To locate significant ITC peaks displayed in Figures 4 and 8A, we first determined the number of ITC peaks across all electrode contacts spanning a brain structure, for each of the 69 frequencies (0.7-50Hz) within the post-event time window (0 to 300ms) using the *findpeaks* function in Matlab. For each trial block, the statistical significance of each ITC peak was evaluated using Rayleigh’s uniformity test. A Bonferroni corrected p-value was determined individually for each frequency based on the number of ITC peaks detected across all electrode contacts within a brain structure at that frequency (i.e. p-value corrected for multiple comparisons = α / number of detected peaks across all electrode contacts at that frequency).

To statistically compare the broadness of significant ITC peak distribution across frequency bands between evoked and modulatory responses (Fig. 6), we quantified the cumulative area under the significant ITC peak distribution curve for each trial block using Matlab’s *cumtrapz* function. Frequency band boundaries were predefined, and the cumulative area between each pair of boundaries was extracted to obtain the area under the curve (AUC) for each frequency band. These were then pooled by response type (i.e., evoked vs. modulatory) across all areas and compared using a Kruskal-Wallis test with Bonferroni correction.

To determine the temporal extent of frequency-band-specific differences in significant ITC peak counts across response types within each area, simultaneous 95% confidence bands were estimated using 1,000 bootstraps for each frequency. Binary significance matrices - defined as time–frequency points where the lower 95% confidence limit of one response type (e.g., evoked ITC) exceeded the upper 95% confidence limit of another (e.g., modulatory ITC) - were then collapsed across predefined frequency bands. For each band, significance values were summed across frequencies and converted into a binary time series, where a value of 1 indicated that at least one frequency bin within the band showed a significant difference at that timepoint. This procedure yielded the time-resolved horizontal lines shown in Figures 7B and 8C, which illustrate when ITC peak counts associated with one response type were significantly greater than those for the other within each frequency band.

## RESULTS

### Evoked multiunit activity (MUA) responses in cortical and subcortical brain structures related to auditory, visual and motor sampling events

Laminar field potential and concomitant multiunit activity (MUA – indexing spiking activity in local neuronal ensembles) profiles were obtained with linear array multicontact electrodes (Fig. 1E) from 66 primary auditory cortex (A1), 31 secondary auditory cortex (Belt), 16 ventral division of the medial geniculate body (MGBv), 34 medial geniculate body (nonlemniscal medial and dorsal divisions, grouped together) (MGB), 11 lateral geniculate nucleus (LGN), 21 medial pulvinar (mPul), 14 lateral pulvinar (Pul) and 14 primary motor cortex sites, across 6 awake macaque monkeys (see Fig. 1A-D for representative histological examples). Neuroelectric activity was recorded under three conditions: 1) during the presentation of a rhythmic (1.6Hz) stream of 50dB, 100ms duration broadband noise (BBN) bursts (auditory stimulation condition), 2) during the presentation of a rhythmic (1.8Hz) stream of LED flashes (visual stimulation condition), and 3) in the absence of any stimuli during which saccades were detected (resting state condition). NHPs were seated in a dim, electrically shielded, sound-attenuated chamber and remained alert, but no behavioral task was required in any condition. Eye position was continuously monitored and the NHPs were free to look around anywhere in the room and at their naturally preferred pace. (see Methods for more details.) One trial block of each condition was recorded in each of the sites. For cortical recordings, CSD profiles were calculated from field potential profiles. CSD is used as an index the location, direction, and density of transmembrane current flow, which is the first-order neuronal response to synaptic input (Nicholson & Freeman, 1975; C E Schroeder, Mehta, & Givre, 1998). Since it is uncertain whether the neural tissue in thalamic areas meets the fundamental assumptions required for applying 1D CSD, which are that neuronal tissue is infinite, homogeneous and an isotropic conductive medium (Mitzdorf, 1985; Nicholson & Freeman, 1975), we opted to calculate the bipolar derivation instead for our thalamic recordings. This method still localizes neural activity and, similar to CSD, is less susceptible to contamination by volume conduction (Biagioni et al., 2025; Kajikawa & Schroeder, 2011).

Figure 1E shows example laminar bipolar field potential and MUA profiles to a BBN stream from a representative MGBv site (top), LGN site (middle) and mPul site (bottom). As expected, responses in MGBv are characteristic of a driving or evoked type consistent with its strong ascending projections from the central nucleus of the inferior colliculus. This response is marked by an early (∼5ms post-stimulus), large-amplitude, and sustained increase in MUA across all recording contacts with an accompanying increase in the bipolar field potential signal (Fig. 1E, top). In contrast, the LGN site shows no suprathreshold post-stimulus firing or appreciable bipolar response to BBN presentation (Fig. 1E, middle). This is not surprising as the LGN is the primary visual nucleus in the thalamus. However, the mPul site, a multisensory thalamic structure, exhibits an intermediate response profile: BBN stimuli evoke an early (∼8ms post-stimulus), transient increase in MUA along with a corresponding bipolar field potential response. However, both signals are lower in amplitude than those observed in MGBv (Fig. 1E, bottom, note different amplitude scales).

When we consider responses to visual stimuli (LED flashes) in these same subcortical recording locations as Figure 1E (Fig. 1F), we find that MGBv shows minimal bipolar or MUA activity. (top). Also unsurprisingly, LGN responds with a large amplitude bipolar field potential and a concomitant sustained MUA increase approximately 100ms post LED flash (Fig. 1F, middle). Akin to the response profile that auditory stimuli elicit in MGBv, this is again characteristic of an evoked type response. The mPul site also shows an evoked response, with suprathreshold MUA activity emerging at ∼120 ms, accompanied by a lower-amplitude bipolar signal relative to LGN (Fig. 1F, bottom, note the different amplitude scales).

In addition to environmental stimuli (i.e. auditory and visual) we also examined whether saccades, an internally generated motor sampling pattern, could result in evoked responses within any of the 8 cortical and subcortical brain structures we examined. Unlike the auditory and visual stimuli which were presented completely isochronously, saccades occurred quasi-rhythmically (see inset presentation rate bar graphs at the top of Fig. 1E-G). The distribution of median saccade rates across all 207 resting state trial blocks illustrates this and also supports similar findings in numerous human and non-human primate studies (Barczak et al., 2019; Hoffman et al., 2013; Maldonado et al., 2008; Rayner, 1998). The median rate of the individual median saccade rates was 2.2Hz (median absolute deviation = 0.24Hz) which is slightly lower than in previous non-human primate studies (Barczak et al., 2019; Berg, Boehnke, Marino, Munoz, & Itti, 2009; Ito, Maldonado, Singer, & Grün, 2011), likely due to the lack of a visual task and strict criteria used for saccade detection (see Methods).

Aligned to saccade onset, the MGBv site again shows minimal bipolar or MUA modulation (Fig. 1G, top). However, in the representative LGN site (Fig. 1G, middle) there is clear MUA suppression prior to and around the time of saccade onset, which is followed by a large amplitude MUA excitation approximately 50ms after saccade onset. As has been documented previously in the LGN, this suppressive/excitatory MUA pattern is typical of saccadic suppression followed by excitation related to the visual input at saccade end (Lee & Malpeli, 1998; Reppas, Usrey, & Reid, 2002). The MUA enhancement is accompanied by a large-amplitude bipolar signal and is comparable in magnitude to the LED-evoked response, although occurring at shorter latency. This is possibly due to the fact that saccading even in a dim room provides richer stimulation than a visually weak (low intensity) stimulus like the LED flash, which would result in slow retinal integration and hence longer latency activation. In the mPul site, saccades do not elicit clear suprathreshold MUA responses but produce a low-amplitude bipolar modulation emerging ∼60 ms after saccade onset (Fig. 1G, bottom).

Next, we wanted to quantify the suprathreshold MUA changes seen in the representative sites in Figure 1, across all trial blocks recorded in the 8 brain regions. To do this we statistically tested whether MUA recorded at any linear array electrode contact within a structure significantly differed in amplitude from a “control” MUA signal, created from random segments of the “real” MUA recorded at the same electrode contact, using Wilcoxon rank sum tests (p < 0.05 with Bonferroni correction) for at least 4ms (see Methods), within a traditional post-stimulus/post-saccadic timeframe (0 to 300ms). If multiple recording contacts within a structure showed significant evoked MUA responses, we selected the contact with the earliest significant response relative to stimulus onset (0ms), whether excitatory or inhibitory, for display in Figure 2.

During the post-stimulus timeframe, we found that many recording locations exhibited BBN-related MUA responses that were significantly larger than their corresponding “control” MUA, which we termed excitatory evoked responses. The top left waveform in Figure 2A displays the grand average of MUA recorded on those recording contacts which displayed excitatory BBN related responses (122/207 or 59% of BBN trial blocks), across all 8 brain structures. A few of the recording locations displayed BBN related MUA responses that were significantly suppressed compared to their matching “control” MUA signal, these we termed inhibitory evoked responses. The waveform in Figure 2A (right) shows the grand average of BBN related inhibitory responses (4/207 or 1.9% of BBN trial blocks). The fact that an obvious post-stimulus MUA suppression is not visible in the grand average is most likely due to the differing latencies of the inhibitory responses across the 4 trial blocks, and thus the responses cancel each other out when averaged. However, the latency of the BBN related off responses are coincident across the 4 trial blocks, and so there is a visible MUA enhancement right after 100ms. The bar graph in Figure 2A displays the distribution of excitatory/inhibitory evoked responses across the 8 subcortical and cortical brain structures. The regional distribution of these responses for BBN stimuli is summarized in columns 1 and 2 of Table 1.

**Table 1.**
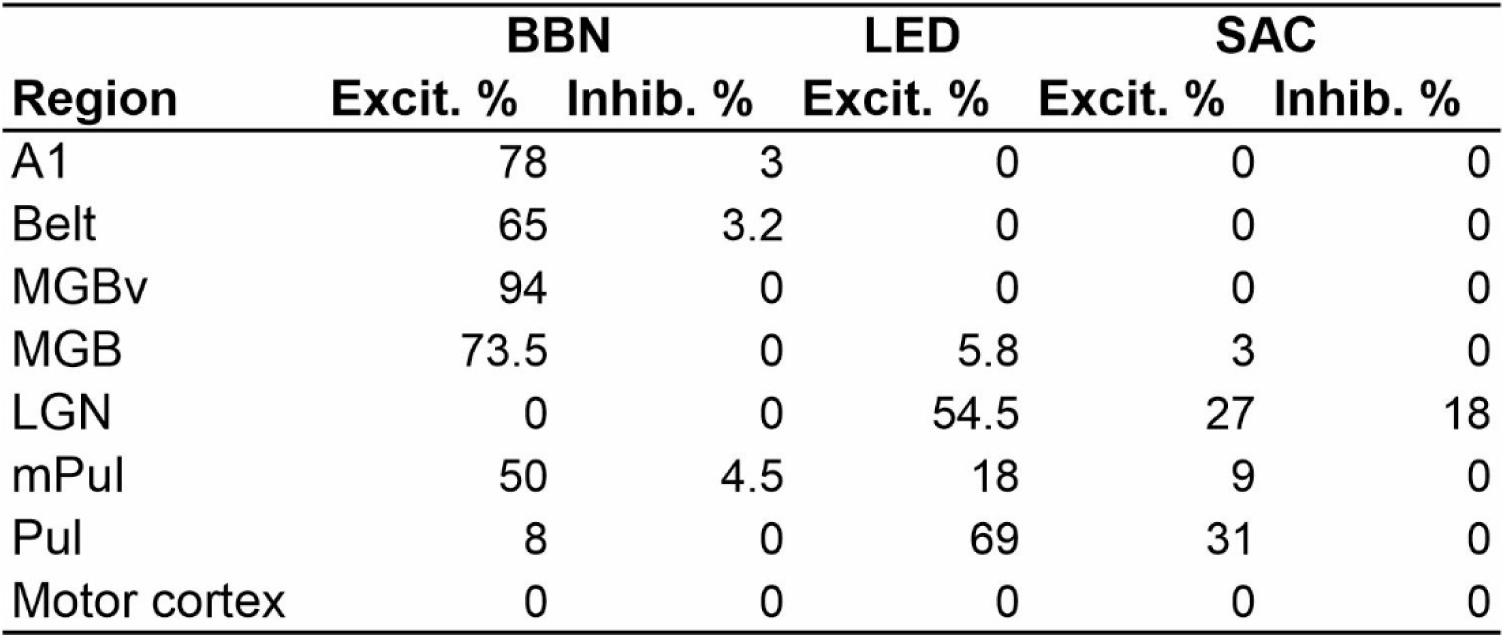
Percent of recording sites showing excitatory (Excit.) or inhibitory (Inhib.) MUA responses across cortical and thalamic regions for auditory (BBN), visual (LED), and saccade (SAC) conditions.

During the post-LED flash timeframe (0-300ms), we found that 21/207 or 10.1% of LED trial blocks showed excitatory MUA evoked responses (see waveform plots in Fig. 2B). None of the brain structures we recorded from exhibited significant inhibitory responses to LED flashes. The distribution of brain structures showing excitatory evoked LED flash related MUA responses is displayed in the mustard bar graph (Figure 2B), the corresponding values are provided in column 3 of Table 1.

When we statistically tested saccade related MUA responses, we found that during the post-saccadic timeframe 10/207 or 4.8% of resting state trial blocks resulted in excitatory saccade related evoked MUA responses (Fig. 2C, left waveform), while only 2/207 or 0.96% of resting state trial blocks resulted in evoked inhibitory responses (Fig. 2C, right waveform). Figure 2C (bar graph) shows the distribution of excitatory and inhibitory evoked MUA responses to saccades across brain structures; the corresponding values are provided in columns 4 and 5 of Table 1.As the brain structures which showed the majority of post-saccadic saccade related excitatory responses are considered traditionally visual or multisensory areas (e.g. LGN and Pul), it is likely that the excitation seen is due to the volley of visual information that enters the brain at saccade end rather than being purely motor related.

### Spectral properties of responses related to auditory, visual and motor sampling events

In the past, a common approach to ascertain whether a brain region was involved in the processing of a certain modality was to test if a stimulus of that modality elicited a suprathreshold evoked response (i.e. neuronal firing) in that area (Hubel & Wiesel, 1962; Mountcastle, 1957; Sherman & Guillery, 2002), not unlike the analysis conducted above. In more recent years it has become clear that stimulus related inputs do not always result in obvious “output like” changes in firing rates of neuronal ensembles. Instead, these inputs can adjust the excitability phase of subthreshold ongoing neuronal oscillatory activity, giving rise to modulatory responses. In sensory cortices, this response type has been shown to be most prevalent for non-preferred modality stimuli (P Lakatos et al., 2007; Peter Lakatos et al., 2008) and saccades (Barczak et al., 2019; O’Connell et al., 2020). Our next set of analyses focused on identifying the signatures of modulatory responses across the 8 brain structures to the different modality events. To do this, we investigated the phase coherence of oscillatory activity across trials (indexed by intertrial coherence - ITC) (P Lakatos et al., 2007; Peter Lakatos, Shah, et al., 2005). The value of ITC will be 1 in the extreme case where the oscillatory excitability phase is the same in each trial, and it will be 0 if the oscillatory excitability phase across trials is random. Time-frequency plots in Figure 3 display average (across all electrode contacts within a structure) auditory, visual and saccade related ITC values derived from the bipolar signal, from the same three representative subcortical structures shown in Figure 1. At the MGBv site, ITC in response to BBN stimuli is marked by high values across nearly all frequencies within the conventional oscillatory frequency bands (delta through gamma, 1-50Hz), consistent with an evoked-type waveform (Lakatos et al., 2007, 2009). In contrast, LED flashes and saccades elicit lower overall ITC values, confined to specific frequencies within select oscillatory frequency bands - such as delta, theta, and alpha for LED, and delta and theta for saccades (Fig. 3A, middle and right). This phenomenon of ITC values confined to restricted frequencies accompanied by no detectable MUA firing (see Fig. 1F & G), is thought to be characteristic of a modulatory response (e.g. Lakatos et al. 2007, 2009). Note that in the delta frequency range, ITC values are higher at the rate of stimuli presentation or saccades (marked by white dotted lines). This is due to the entrainment or alignment of the neuronal oscillatory fluctuations (with a similar excitability phase) to the rhythmically presented stimuli and quasi-rhythmic saccades.

**Figure 3.**
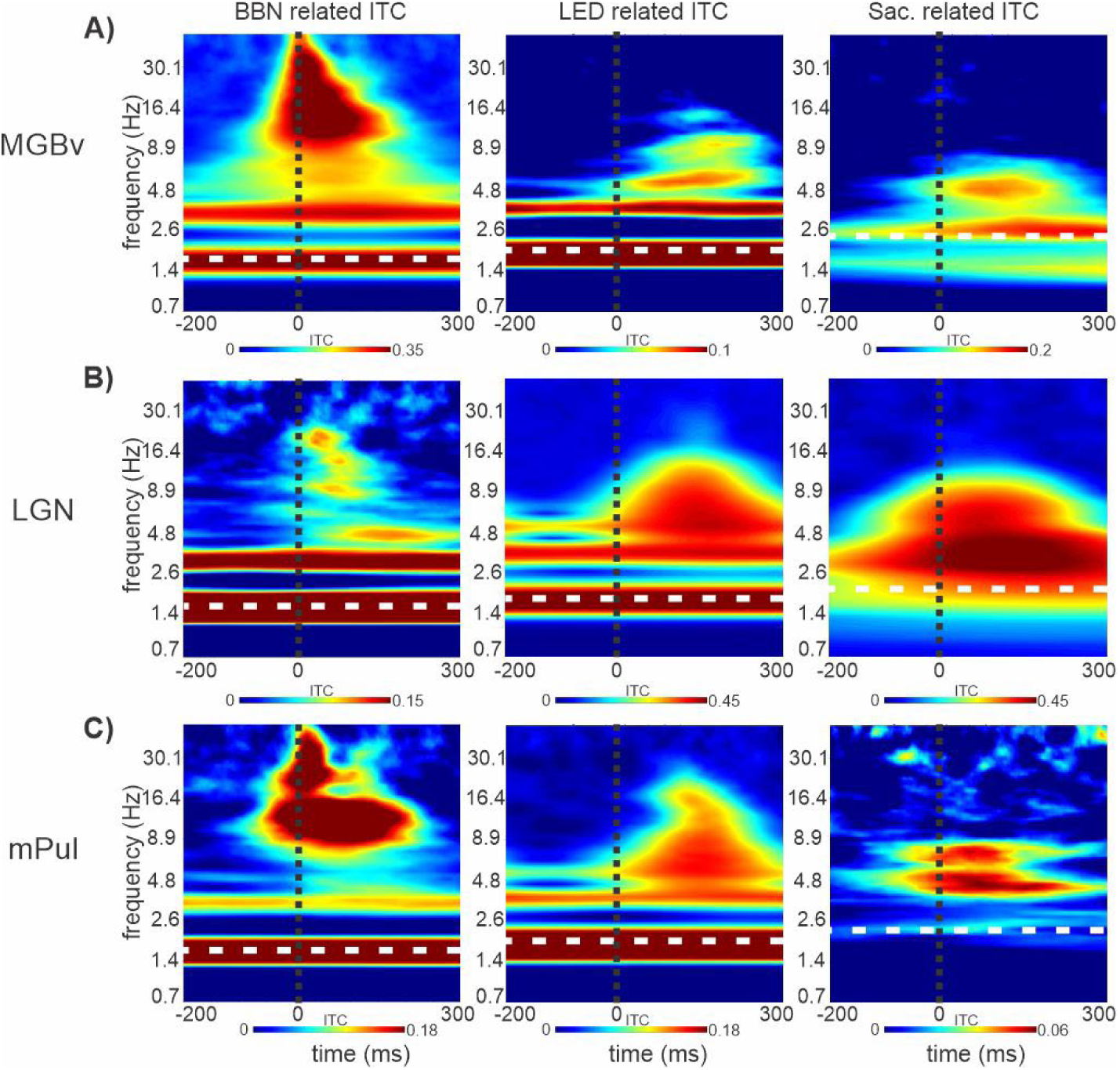
Spectrotemporal properties of event related responses to preferred and non-preferred modality events in subcortical structures. (**A–C**) Time–frequency plots of bipolar inter-trial coherence (ITC) for representative recordings from MGB (**A**), LGN (**B**), and medial pulvinar (mPul; **C**), corresponding to the sites shown in Fig. 1. White dotted horizontal lines indicate the stimulus or event rates (BBN: 1.6 Hz; LED: 1.8 Hz; saccades: median rate for each block (**A**: 2.4 Hz; **B**: 2.17 Hz; **C**: 2.32 Hz), and black vertical lines mark time 0ms.

In the representative LGN site, LED flashes and saccades resulted in high ITC values broadly across delta through alpha frequency bands (Fig. 3B), indicating the evoked responses seen in Figure 1. Since there are no known auditory inputs to LGN and no observed evoked response (Fig. 1E), the reduced ITC values peaking at specific frequencies within the delta (approximately 1.6Hz - implying entrainment), theta, alpha and beta frequency bands in response to the BBN stimuli is suggestive of a modulatory response.

In the multisensory mPul site, although ITC values were lower overall, BBN and LED flashes lead to significant ITC values at the delta stimulation rates, indictive of neural entrainment. This was coupled with broadband ITC values across a range of frequency bands, consistent with the evoked responses seen in Figure 1E & F (Fig. 3C). In contrast, saccadic events produced much lower ITC values, which were almost exclusively restricted to the theta frequency band.Hz in some brain regions could reflect a harmonic of the true saccadic rhythm rather than direct entrainment.

To determine which sites exhibited a purely modulatory (signified by clustering of significant ITC values at specific frequencies) versus an evoked (indicated by significant broadband ITC across multiple frequency bands paired with a suprathreshold MUA) response, we first searched for significant ITC peaks. ITC peaks were evaluated using Rayleigh’s uniformity test (p < 0.05 with Bonferroni correction), across individual recording contacts within the post-event time windows (i.e. 0 to 300ms) and across each frequency band (1-50Hz) during all trial blocks (see Methods). Traces in Figure 4A show the frequency distribution of significant BBN related ITC peaks (normalized to the peak of each trace for visualization purposes) obtained from BBN trial blocks which resulted in evoked MUA responses (brown traces) vs non-evoked (green traces), across the 8 brain structures. Counts of significant BBN-related ITC peaks for each region are shown in the first two columns of Table 2. A prominent ITC peak at the BBN presentation rate of 1.6 Hz (red dotted line) was observed across all brain regions, particularly for non-evoked trial blocks, indicating strong entrainment of local delta oscillations to a consistent phase by the BBN stream. In regions where BBN evoked robust MUA responses - such as A1, Belt, and MGBv (see Fig. 2A), ITC responses also showed a broad distribution, with a “tail” of an equally large number of ITC peaks across the conventional oscillatory frequency bands. In contrast, trial blocks without evoked responses (e.g., in the LGN and motor cortex) displayed ITC peaks that were more narrowly restricted to distinct frequency bands. This pattern suggests that BBN stimuli organize ongoing oscillatory activity on distinct timescales, as evidenced by the frequency-specific distribution of ITC peaks.

**Figure 4.**
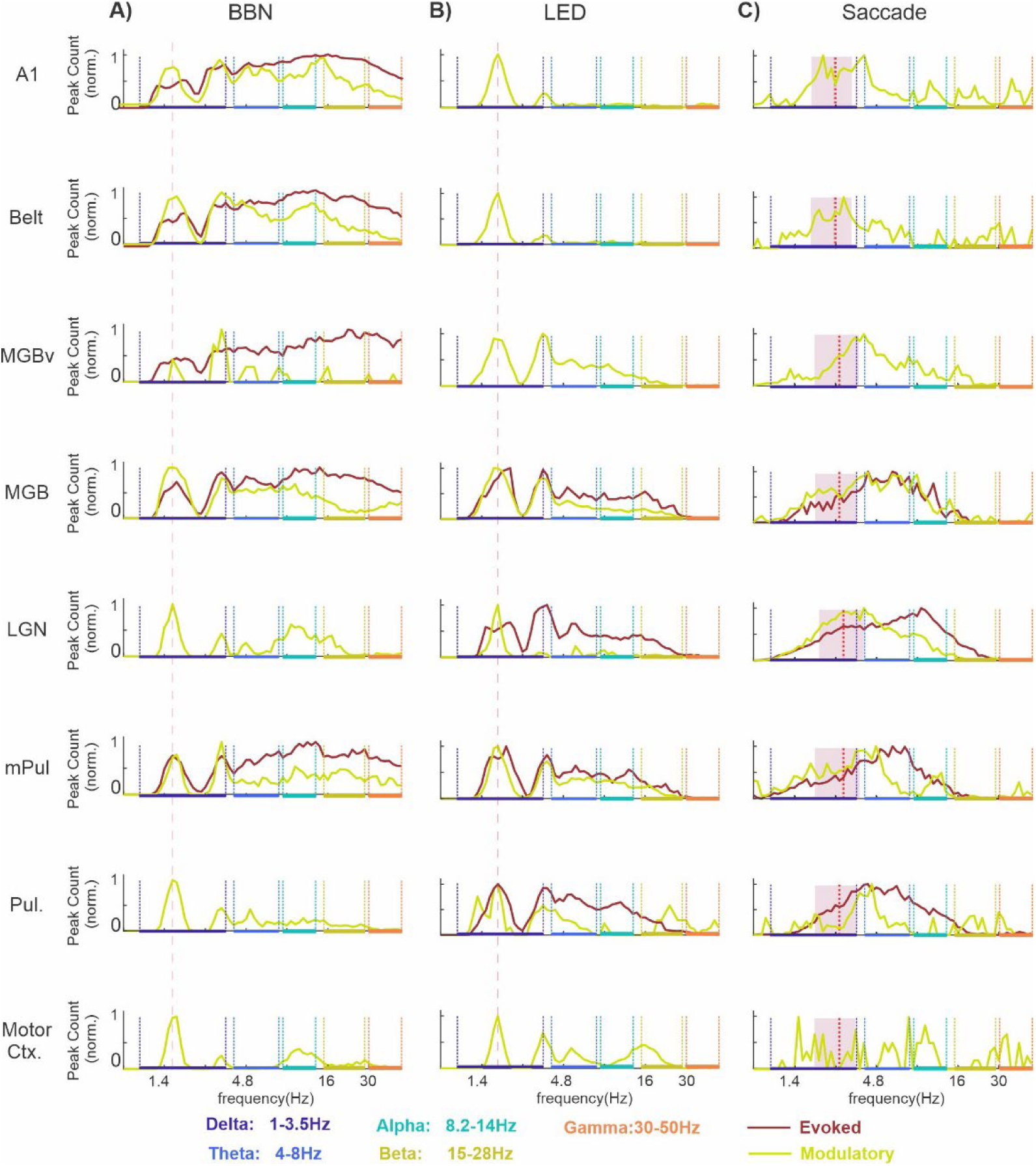
Frequency distribution of significant CSD and bipolar ITC peaks related to multi-modal events across brain structures. (**A–C**) Frequency distributions of significant ITC peaks (1–50 Hz, 0–300 ms) across cortical and subcortical regions. Traces show evoked responses (brown) and non-evoked (modulatory) responses (green). Colored bars on the x-axis indicate the five frequency bands used for subsequent analyses. **A**) BBN-related responses; the red vertical line indicates the BBN presentation rate (1.6 Hz). **B**) LED-related responses, shown with the same format as in A. **C**) Saccade-related responses; red vertical lines indicate median saccade rates for each structure, and mauve shading denotes +/-median absolute deviation.

**Table 2.** Number of significant ITC peaks across regions and conditions (BBN, LED, SAC), separated into evoked and modulatory (Modul.) responses; blanks indicate no significant peaks.

|  | BBN |  | LED |  | SAC |  | BBN | LED | SAC |
| --- | --- | --- | --- | --- | --- | --- | --- | --- | --- |
| Region | Evoked | Modul. | Evoked | Modul. | Evoked | Modul. | Total | Total | Total |
| A1 | 74231 | 4961 |  | 10343 |  | 1034 | 79192 | 10343 | 1034 |
| Belt | 24452 | 5410 |  | 5082 |  | 873 | 29862 | 5082 | 873 |
| MGBv | 8930 | 44 |  | 4735 |  | 1670 | 8974 | 4735 | 1670 |
| MGB | 12823 | 2715 | 2362 | 13436 | 672 | 4142 | 15538 | 15798 | 4814 |
| LGN |  | 652 | 11092 | 296 | 6696 | 2515 | 652 | 11388 | 9211 |
| mPul | 11031 | 1663 | 3891 | 9766 | 1433 | 1408 | 12694 | 13657 | 2841 |
| Pul |  | 1599 | 9761 | 960 | 3231 | 490 | 1599 | 10721 | 3721 |
| Motor Crtx. |  | 654 |  | 2449 |  | 80 | 654 | 2449 | 80 |

Although the total number of significant LED-related ITC peaks was lower than for BBN stimuli, the overall pattern across regions was similar (see Table 2, columns 3-4). Every sampled brain structure showed a substantial number of significant ITC peaks at the rate of LED flash presentation (1.8 Hz; red line), indicating entrainment. This effect was particularly pronounced in areas in which non-evoked responses to LED flashes predominated (e.g., green traces Fig. 4B). Additionally, in recording sites that exhibited primarily evoked responses to LED stimuli - such as the LGN and Pul (e.g., brown traces in Fig. 4B) - significant ITC peaks were broadly distributed across adjacent frequency bands, ranging from delta through beta. In the case of non-evoked responses, ITC peaks were typically concentrated within the delta band near the stimulation rate (e.g., A1 and Belt), or clustered around distinct frequencies within specific bands, such as in the Motor Cortex. Unlike BBN stimuli, LED flashes did not elicit many significant ITC peaks in the gamma frequency band. Notably, a prominent number of ITC peaks related to both BBN and LED flash stimuli were observed around 3 Hz across all recording sites. This likely reflects a harmonic relationship, as 3 Hz is a shared harmonic of the BBN and LED presentation rates (1.6 Hz and 1.8 Hz, respectively).

Saccade-related responses yielded the lowest number of significant ITC peaks across regions; detailed counts are provided in the fifth and sixth columns of Table 2. Surprisingly, the ITC peak distribution traces shown in Figure 4C appear relatively ‘flatter,’ lacking distinct, sharply defined peaks. For non-evoked responses, ITC peaks are primarily concentrated in the high delta to lower theta frequency range. In contrast, saccade-related evoked responses are biased toward higher theta to alpha frequencies, with ITC peaks predominantly concentrated within these bands. A clear distinction between the ITC frequency distributions for sensory stimuli and those related to saccades is the absence of a prominent peak at the median saccade rate (indicated by red lines), which would typically suggest entrainment to a rhythmic saccadic pattern. One possible explanation is that our strict criteria for saccade detection (see Methods) may have led to the omission of some saccades, resulting in an underestimated median rate. Alternatively, if all saccades were accurately detected, the observed peak around 4 Hz in some brain regions could reflect a harmonic of the true saccadic rhythm rather than direct entrainment.

A summary of the results thus far is presented in Figure 5. The height of each bar shows the percentage of trial blocks recorded in each of the 8 brain structures that displayed significant ITC in at least one frequency band and at least one linear array electrode contact, while the height of the brown area within the bar illustrates what percentage of those trial blocks also exhibited a concurrent significant MUA response. In other words, the brown areas denote prevalence of evoked type responses (same data as displayed in bar graphs in Fig. 2), while the green designates the incidence of modulatory type responses - clustering of significant ITC values at specific frequencies without an accompanying suprathreshold MUA response. In the case of the BBN streams, trial blocks in nearly all areas (with the exception of mPul) exhibited predominantly either evoked or modulatory response profiles (Fig. 5A), with the regional distribution summarized in columns 1 and 2 of Table 3.Visual stimulation likewise affected a large proportion of trial blocks in each of the 8 brain structures (Fig. 5B), with the corresponding distribution shown in columns 3 and 4 of Table 3. Likewise, during resting-state trial blocks, saccades elicited evoked and modulatory responses across regions (Fig. 5C), as summarized in columns 5 and 6 of Table 3. Clearly auditory and visual stimuli, even when they do not match the preferred modality of a specific brain region, absolutely impact neuronal excitability even at a subthreshold level, as evidenced by the total height of each bar approaching 100% in Figure 5A and B. It is also apparent from these bar graphs which of the 8 brain structures are part of the auditory or visual driving pathway that transmits detailed stimulus specific information (i.e. bars with taller brown regions), and which belong to the modulatory pathway that alters subthreshold neuronal excitability (i.e. bars with taller green regions). Similarly, inputs related to motor sampling patterns (i.e. saccades) are also capable of impacting neuronal excitability at both sub- and suprathreshold levels as demonstrated by the tallness of the bars in Figure 5C. While it is probable that the modulation seen in visual areas (e.g. LGN and Pul) is due to visual input at saccade end rather than being motor related, a previous finding that saccades and visual stimuli reset delta/theta oscillations to opposing excitability phases (O’Connell et al., 2020) suggests that this is unlikely in non-visual brain regions (e.g. A1, MGBv, etc.). Therefore, it is apparent that the motor system has an impact on sensory processing at all hierarchical levels mainly through the reorganization of ongoing neuronal oscillations.

**Figure 5.**
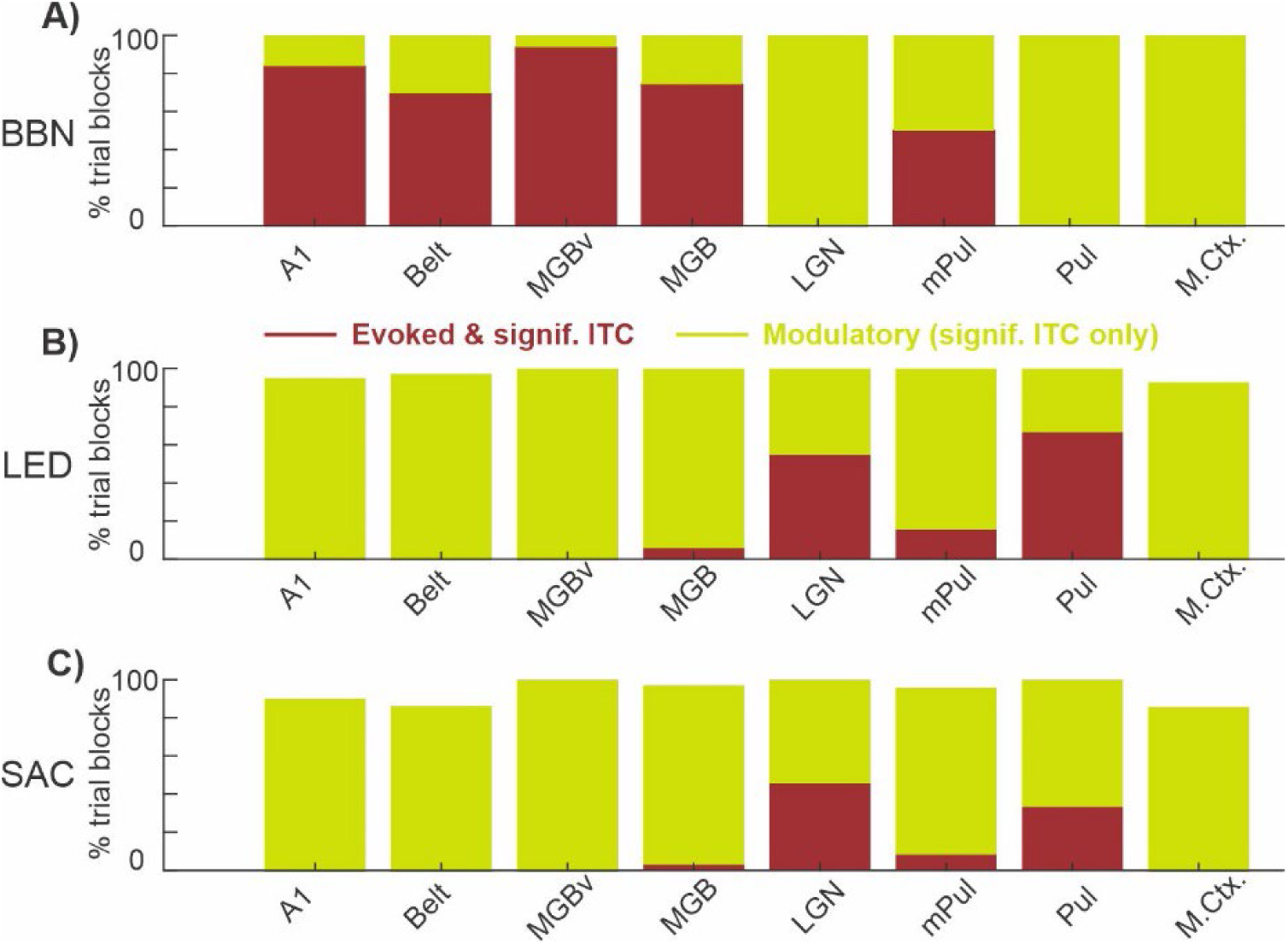
Prevalence of evoked and modulatory responses related to multi-modal events across brain structures. **A–C**) Bar graphs show the percentage of trial blocks within each brain structure exhibiting significant ITC responses (0-300ms) for each condition. Brown segments indicate evoked responses (significant ITC accompanied by significant MUA), while green segments indicate modulatory responses (significant ITC without concurrent MUA). **A**) BBN-related responses. **B**) LED-related responses. **C**) Saccade-related responses.

**Table 3.**
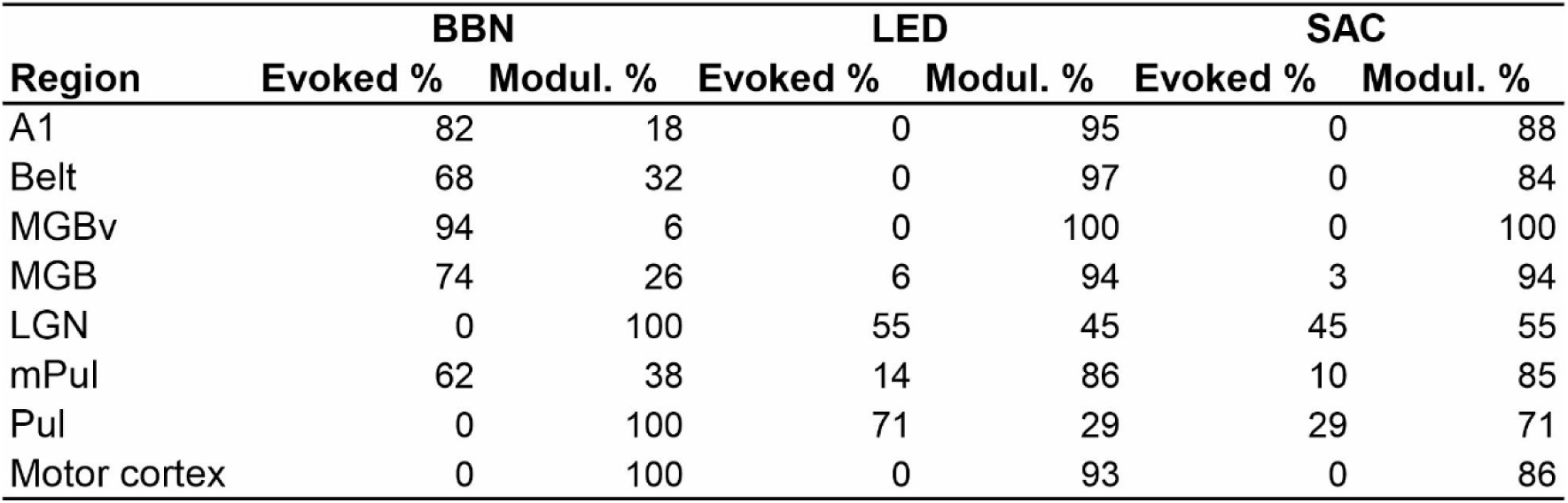
Percent of sites with evoked or modulatory (Modul.) responses across regions and conditions (BBN, LED, SAC).

### Evoked and modulatory responses exhibit distinct AUC patterns across adjacent frequency bands

Given that trial blocks with evoked MUA responses exhibited a qualitatively broader distribution of significant ITC peaks compared to modulatory blocks we sought to quantify how this difference was expressed across neighboring frequency bands. To do so, we computed the area under the curve (AUC) for the frequency distribution of significant ITC peaks within five canonical frequency bands (see colored bars on the x-axes in Fig. 4) both response types. We hypothesized that evoked responses would yield relatively uniform AUCs across adjacent bands, whereas modulatory responses would show disproportionate increases in specific bands, resulting in greater differences between neighboring frequency bands. To test this, we pooled AUCs within each frequency band by response type (evoked vs. modulatory) across all areas and conducted statistical comparisons across frequency bands. This analysis was performed separately for each event type (Fig. 6). For evoked responses to BBN streams, AUCs were largely comparable across neighboring frequency bands, with only a modest deviation in the beta band, which differed significantly from its adjacent alpha and gamma bands (Kruskal–Wallis test with Bonferroni correction, p < 0.01; Fig. 6A, top). In contrast, BBN-related modulatory responses showed pronounced differences across adjacent bands, with delta band AUCs significantly exceeding those of all other frequency bands, including the neighboring theta band (p < 3.41 × 10⁻⁷; delta vs. theta: p = 3 × 10⁻⁸; Fig. 6A, bottom). For LED-related evoked responses, delta band AUCs were significantly greater than those of higher-frequency bands (alpha - gamma; p < 0.001), but did not differ from the adjacent theta band (p = 0.2), indicating relatively uniform AUCs across neighboring bands (Fig. 6B, top). In contrast, LED-related modulatory responses exhibited strong band-specific effects, with delta band AUCs significantly larger than those of all other frequency bands, including the adjacent theta band (p < 1 × 10⁻³⁰; delta vs. theta: p = 1 × 10⁻³¹; Fig. 6B, bottom). These statistical comparisons in the context of external sensory stimuli support our hypothesis that evoked responses are characterized by relatively uniform AUCs across adjacent frequency bands, whereas modulatory responses show marked band-specific concentrations.

**Figure 6.**
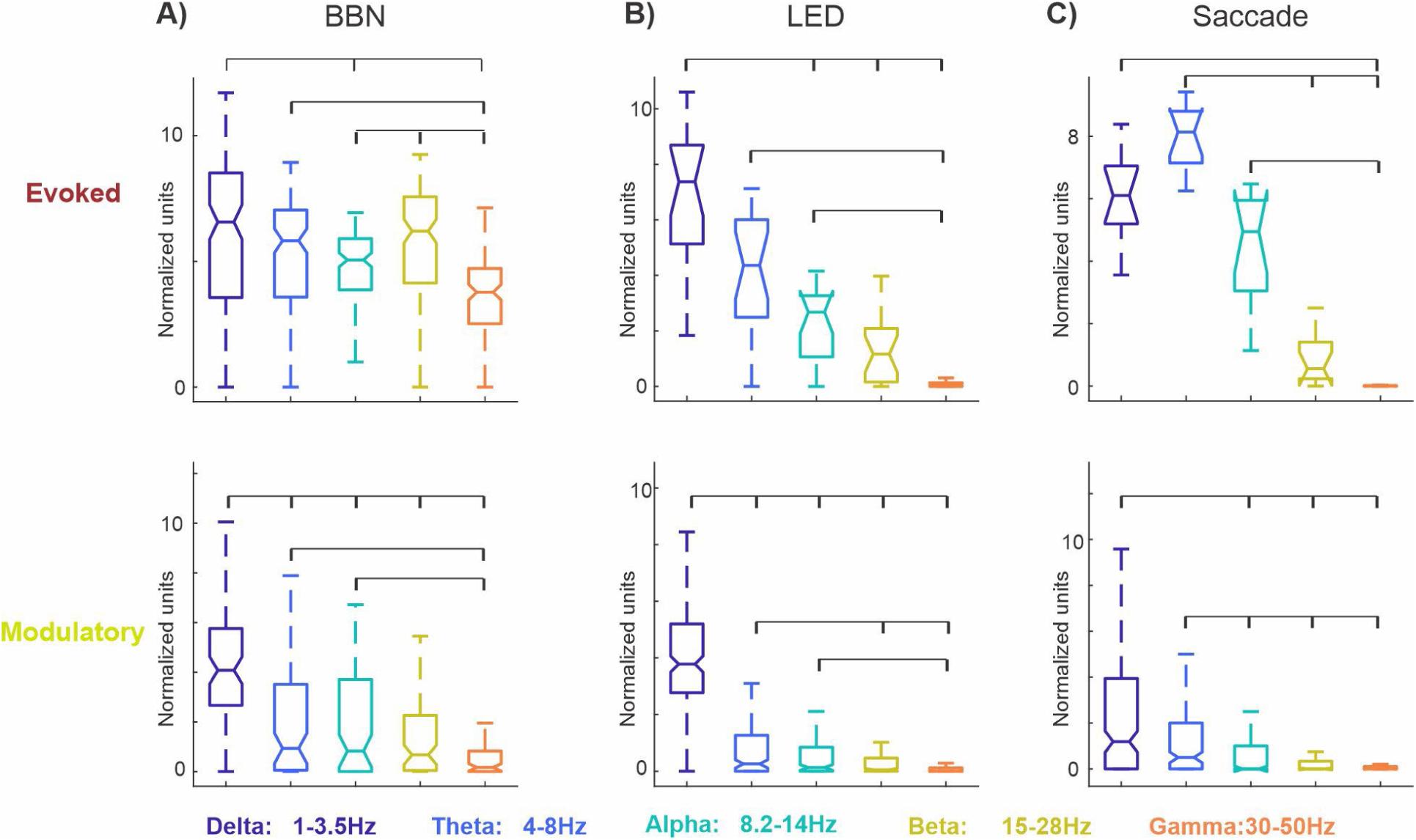
Statistical comparison of frequency band-limited AUCs of significant ITC peak distributions. **(A–C)** Boxplots show pooled area under the curve (AUC) values for the frequency distribution of significant ITC peaks across five canonical frequency bands, separated by response type (evoked, top; modulatory, bottom) and condition. AUCs were computed from trial blocks exhibiting evoked or modulatory responses and pooled across all brain regions. Brackets indicate significant differences between frequency bands (Kruskal–Wallis test with Bonferroni correction, p < 0.01). **A)** BBN-related responses. **B)** LED-related responses. **C)** Saccade-related responses.

For saccade-related evoked responses, although theta band AUCs were the largest, they differed significantly only from higher-frequency bands (beta and gamma; p < 1.2 × 10⁻³⁴; Fig. 6C, top) and not from the adjacent alpha band. This pattern is consistent with that observed for BBN- and LED-related evoked responses, in which AUCs are relatively similar across neighboring frequency bands. While saccade-related modulatory responses resembled sensory stimulus–related modulatory responses in exhibiting the largest AUCs in the delta band (Fig. 6C, bottom), these AUCs were not significantly greater than those in the adjacent theta band (p = 0.034). This indicates a broader distribution of ITC peaks across neighboring frequency bands, more similar to the pattern observed for evoked responses than for sensory-driven modulatory responses. Together, these results suggest that phase-resetting signatures associated with internally generated motor events (e.g., saccades) are less frequency-specific than those driven by external sensory stimuli, possibly due to differences in the underlying circuitry.

### Spectrotemporal distribution of significant ITC peaks related to multi-modal events across brain structures

Up to this point, our analysis has focused on the distribution of significant ITC peaks pooled across the entire 0-300ms post-stimulus period. To gain deeper insight into the temporal dynamics of these responses, we next examined their joint distribution across time and frequency (i.e., their spectrotemporal profile). Figure 7A shows color plots of the mean spectrotemporal distribution of significant ITC peaks within each brain area. In generating this figure, BBN or LED trial blocks that elicited evoked MUA responses were used for regions associated with auditory or visual driving pathways, respectively (i.e., areas with taller brown bars in Fig. 5A–B). If no evoked responses were evident, then BBN, LED and saccade modulatory trial blocks were used to construct the color plots. From visual inspection, several features are notable. First, consistent with the Heisenberg uncertainty principle (Cohen, 2020), delta-band ITC peaks are temporally diffuse, spanning the entire 0–300ms window across all regions. Second, within each area, the timing of ITC peaks varies by event type. For example, in the LGN, LED-related theta/alpha ITC peaks occur ∼50ms later than those associated with saccades, while BBN-related peaks are less frequent and cluster in the alpha band around 100ms following the stimulus. Third, in BBN trial blocks that elicited evoked MUA responses (highlighted by red boxes), gamma ITC peaks appear in two bursts separated by ∼100ms, likely reflecting transient “on” and “off” MUA responses to the 100ms BBN stimulus. These gamma bursts are absent in BBN modulatory trial blocks. Finally, saccade-related ITC peaks are predominantly restricted to the delta–theta frequency range across all regions (cf. Fig. 4C).

**Figure 7.**
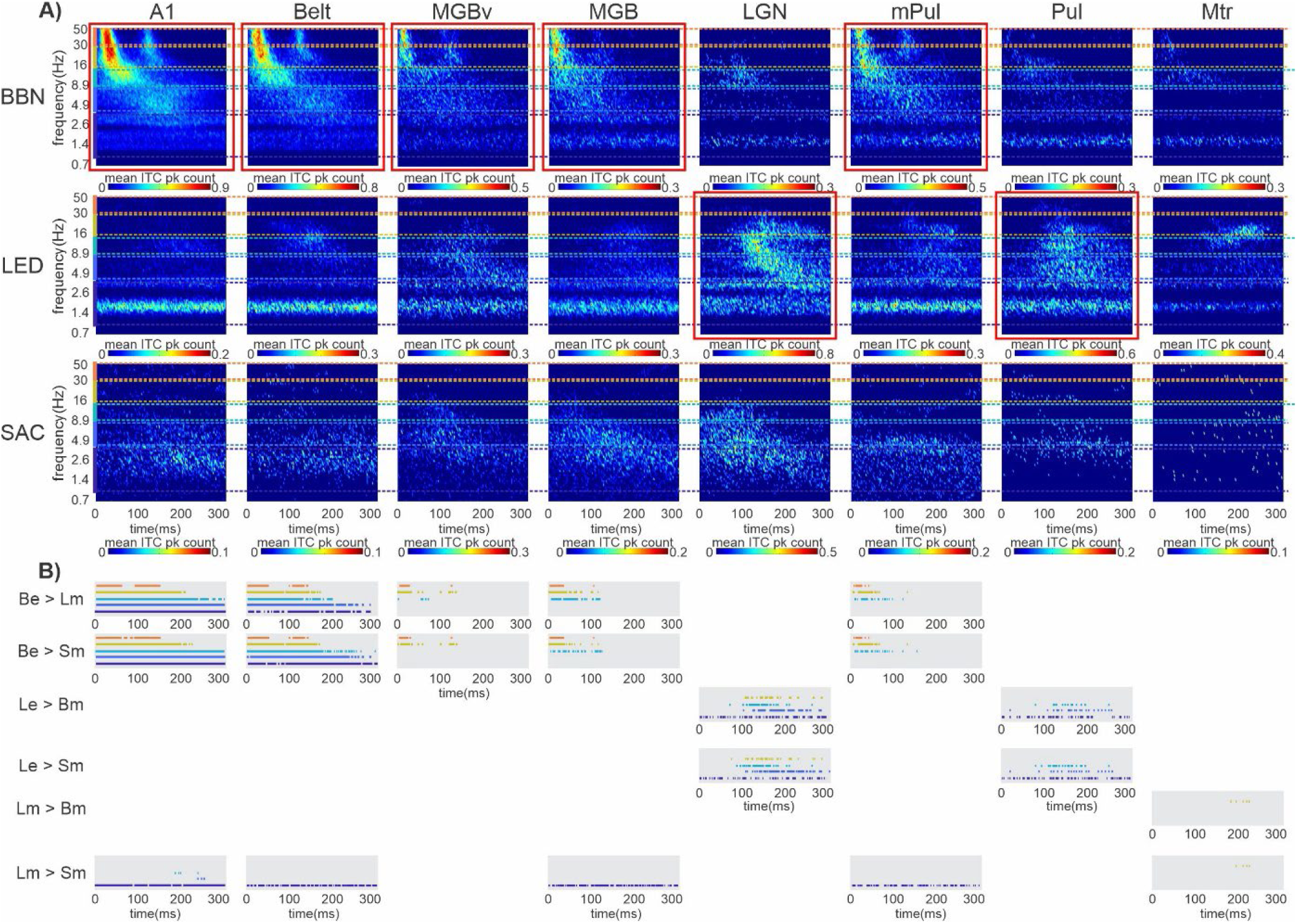
Spectrotemporal distribution of significant CSD and bipolar ITC peaks across brain structures. **A)** Color plots show the mean spectrotemporal distribution of significant ITC peaks across eight brain regions (columns) for three event types (rows). Red boxes indicate plots derived from trial blocks with evoked MUA responses; unboxed plots reflect modulatory trial blocks. Colored bars on the y-axes denote the five frequency bands (see Figs. 4 and 6). **B)** Horizontal lines indicate time intervals where bootstrapped confidence intervals (p < 0.05) for ITC peaks differed between event types. Line color denotes the frequency band (see Figs. 4 and 6). Labels indicate comparison types: Be (BBN evoked), Bm (BBN modulatory), Le (LED evoked), Lm (LED modulatory), and Sm (saccade-related modulatory). Absence of a gray rectangle indicates no significant differences for that comparison within a given region.

To statistically compare the timing of band limited ITC peaks across the three event types within each area, we calculated simultaneous confidence bands of ITC peak count related to each event type (see Methods) and determined the time intervals in which the bands did not overlap (Fig. 7B). As expected, in A1 (Fig. 7B, first column), BBN trial blocks that elicited evoked responses contained significantly more delta-, theta-, and alpha-band ITC peaks than LED and saccade modulatory trial blocks across the entire 0-300ms window i.e., confidence bands did not overlap. In contrast, beta-band ITC peaks were elevated only within the 0-200ms interval, while gamma-band ITC peaks were elevated within 0-60ms and 100-150ms, consistent with transient the ‘on’ and ‘off’ MUA responses to the 100ms BBN stimulus. LED modulatory trial blocks exhibited significantly more delta-band ITC peaks than saccade modulatory trial blocks across the entire 0-300ms window, likely reflecting the delta-rate presentation of the LED stream. Theta- and alpha-band LED-related ITC peaks were significantly greater only at later time points, around 250ms and 200ms respectively, consistent with delayed visual influences on neural phase coherence in A1. In Belt (Fig. 7B, second column), a similar overall pattern was observed.

In the subcortical regions identified as part of the auditory driving pathway (MGBv, MGB, mPul; cf. Fig. 5A) greater ITC peak counts in evoked BBN trial blocks compared to LED and saccade modulatory trial blocks were confined to the higher frequency bands (Fig. 7B, third, fourth, and sixth columns). In MGBv, beta- and gamma-band ITC peaks were elevated within the 0-40ms interval and again around 120ms following stimulus offset. In MGB and mPul, BBN-evoked alpha-band ITC peak counts were also increased relative to LED- and saccade-related modulatory trial blocks during the 0-100ms duration of the BBN stimulus. In contrast, delta/theta ITC peak counts did not differ between BBN-evoked and LED- or saccade-related modulatory trial blocks, indicating that visual and motor events produce a similar number and temporal distribution of low-frequency ITC peaks as the BBN stimulus. This selective enhancement of higher-frequency ITC peaks in BBN-evoked responses, coupled with comparable low-frequency activity across conditions, suggests a distinction between subcortical and cortical auditory driving pathways.

In the main visual thalamic nucleus, LGN, LED trial blocks that elicited evoked responses contained significantly more delta-band ITC peaks than BBN and saccade modulatory trial blocks across the entire 0-300ms window (Fig. 7B, fifth column), likely reflecting the delta-rate presentation of the LED stream. In addition, theta-, alpha-, and beta-band ITC peaks were elevated during the 100-250ms, 90-200ms, and 100-220ms intervals, respectively, consistent with the delayed onset of LED-evoked responses (cf. Fig. 1B). Although saccade- and BBN-related modulatory trial blocks showed clustering of theta- and alpha-band ITC peaks, respectively, around ∼100ms, these peak counts did not differ significantly from those observed in LED-evoked trial blocks. A similar pattern was observed in Pul (Fig. 7B, seventh column), with the exception that beta-band ITC peaks were not increased in LED-evoked trials compared to BBN and saccade modulatory trials. This difference highlights a distinction between LGN and Pul, despite both structures exhibiting robust visual responses and belonging to the visual driving pathway (cf. Fig. 5B), suggesting region-specific differences in how visual thalamic nuclei encode stimulus-driven activity across frequency bands.

In Motor cortex, LED modulatory trial blocks contained significantly more beta-band ITC peaks than BBN and saccade modulatory trial blocks within the 190-225ms (Fig. 7B, last column). No significant differences were observed across delta, theta, alpha, or gamma bands. Across all regions, there were no instances in which saccade-related modulatory ITC peak counts exceeded those observed for LED- or BBN-related modulatory trial blocks.

### Stimulation-rate dependent modulatory phase coherence across frequency bands

The present findings demonstrate that, for modulatory responses, the largest proportion of significant ITC peaks is concentrated in the delta frequency band for BBN and LED stimuli, and in the delta–theta range for saccades (cf. Figs. 4 and 7), consistent with neural oscillatory entrainment to multimodal events. To more directly test whether modulatory entrainment can be driven beyond the delta range, rhythmic streams of LED flashes were presented at a theta stimulation rate of 6 Hz in a subset of recordings (n = 33; 5 A1, 4 Belt, 4 MGBv, 7 MGB, 7 mPul, and 6 Motor cortex sites). Traces in Figure 8A show the frequency distribution of significant ITC peaks related to the 6Hz LED stream (dark green trace), overlaid with the corresponding distribution obtained from the same sites during the 1.8Hz LED stream (light green trace). For visualization purposes, both distributions were normalized to the peak of each trace. Similar to the 1.8Hz condition, the distribution associated with the 6Hz stream exhibits a pronounced cluster of significant ITC peaks at the stimulation rate (purple solid line), consistent with entrainment. A secondary concentration of peaks is evident near 12Hz, likely reflecting a harmonic relationship, with additional grouping in the beta range around 18Hz. As with the 1.8Hz LED stream, few ITC peaks were observed in the gamma band. Notably, the 6Hz condition revealed no ITC peaks in the delta band, indicating that phase resetting at oscillatory frequencies lower than the stimulation rate does not appear to occur.

**Figure 8.**
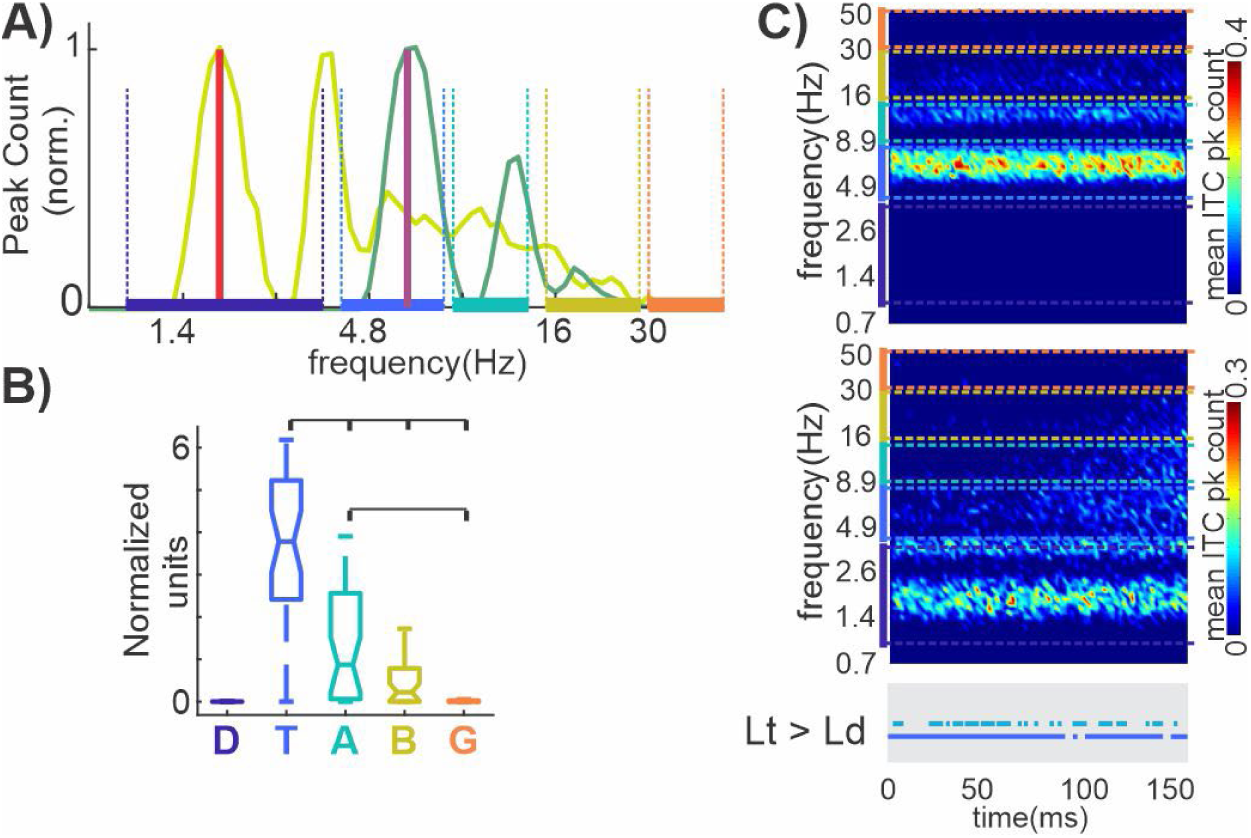
Frequency and spectrotemporal distribution of ITC peaks during theta-rate (6 Hz) LED stimulation. **A)** Frequency distributions of significant ITC peaks for modulatory LED trials at 6 Hz (dark green) and 1.8 Hz (light green) (n = 33 blocks, six regions), computed over 1-50 Hz and 0-150 ms. Vertical lines indicate stimulation rates; colored bars denote frequency bands (see Figs. 4, 6). **B)** Boxplots of AUC values across frequency bands for 6 Hz modulatory trials; brackets indicate significant differences (Kruskal–Wallis with Bonferroni, p < 0.001). **C)** Spectrotemporal distributions for 6 Hz (top) and 1.8 Hz (bottom). Horizontal lines indicate time intervals where bootstrapped confidence intervals differ between conditions (p < 0.05).

Next, we examined whether the frequency signatures of modulatory phase resetting elicited under theta rate stimulation differed from that observed during modulatory responses to the delta rate LED stream within these six brain structures (cf. Fig. 6). As shown in Figure 8B, theta-band AUCs associated with the 6Hz LED related modulatory responses were significantly greater than those of all other frequency bands, including the adjacent alpha band (p < 0.001, Kruskal-Wallis test with Bonferroni correction). These results indicate that modulatory responses, irrespective of the stimulation rate by which they are generated, exhibit a preferential concentration of ITC peaks within a specific frequency band.

Color plots in Figure 8C show the mean spectrotemporal distribution of significant ITC peaks constructed form the same modulatory 6Hz (top) and 1.8Hz (bottom) LED trial blocks. Note that due to the rapid presentation of the 6Hz LED flashes (SOA=166ms) the timescale of these plots is half that of those in Figure 7. Again, it is apparent that there were no ITC peaks observed in the delta range during theta-rate LED stimulation. As expected, theta-rate trial blocks exhibited significantly more theta-band ITC peaks than delta-rate blocks across the entire 0-150ms window (blue horizontal line within grey box), reflecting the rhythmic theta presentation rate of the LED stream. In addition, alpha-band ITC peaks were significantly greater, especially around 20-70ms post LED flash (green horizontal line within grey box), indicating that theta-rate LED stimulation induces earlier phase coherence in both theta- and alpha-frequency bands compared to delta-rate stimulation.

### Phase alignment of MUA across brain structures related to multi-modal events

Results so far show that neuronal oscillations entrain with a similar excitability phase, as indexed by significant ITC values, to the rate of the rhythmically presented external stimuli and quasi-rhythmic saccades. Our next question centered around whether oscillations were being synchronized to a similar high or low excitability phase across areas. Such synchronization could be behaviorally relevant as a means to perhaps facilitate or impede the transfer of sensory information. To investigate this, since MUA has a straightforward relationship with excitability ((Peter Lakatos et al., 2013; O’Connell et al., 2020; O’Connell, Falchier, McGinnis, Schroeder, & Lakatos, 2011), we calculated the mean MUA phase at the frequency of peak ITC values. This was done within the delta band for the BBN and LED stimulus streams, and within the delta-theta band for saccade-related activity. Figure 9A shows stacked colored histograms depicting the distribution of mean translaminar and transnuclei MUA phases for each recording site at event onset. In generating this figure, BBN or LED trial blocks that elicited evoked MUA responses were used for brain structures considered part of the auditory or visual driving pathways (i.e. areas with taller brown bars in Fig. 5A–B) respectively. Otherwise, BBN, LED and saccade modulatory trial blocks were used to construct the phase histograms. Colored asterisks indicate brain areas exhibiting significant phase bias, as determined by Rayleigh’s test for non-uniformity (p < 0.01).

**Figure 9.**
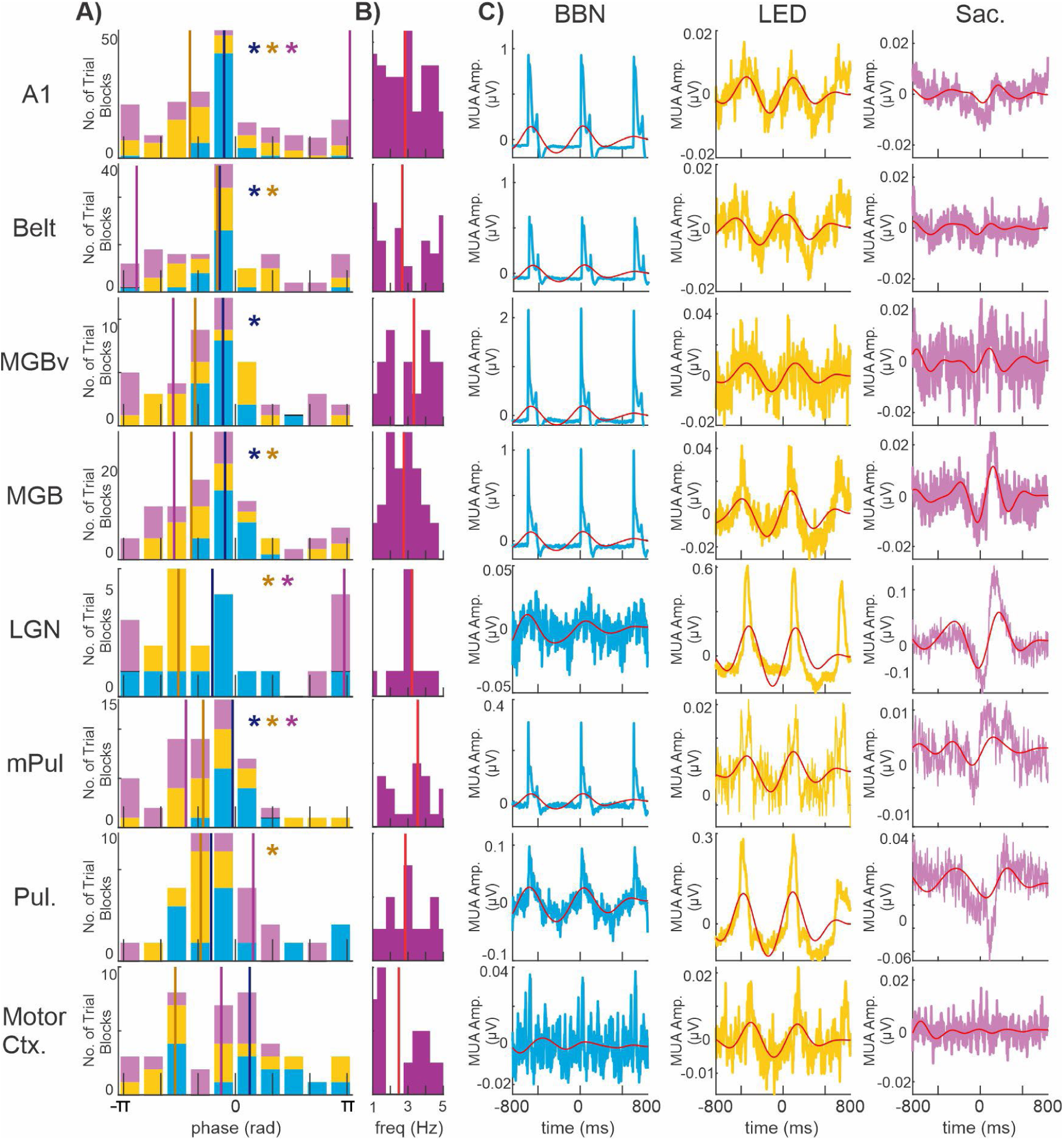
Phase organization of low-frequency MUA activity at the onset of multimodal events across brain structures. **A)** at event onset for BBN bursts (blue), LED flashes (yellow), and saccades (pink), pooled across recording sites within each structure. Phases were computed at the frequency of peak ITC within the delta band for BBN and LED, and within the delta–theta range for saccades. Vertical lines indicate circular mean phase (color-coded by event type). Colored asterisks denote significant phase bias (Rayleigh’s test, p < 0.05). **B)** Histograms show the distribution of peak ITC frequencies derived from MUA signals aligned to saccade onset (0 ms), pooled across sites. The red vertical line indicates the median frequency. **C)** Event-aligned MUA traces for BBN (blue), LED (yellow), and saccades (pink), averaged across trial blocks within each structure. Red traces show bandpass-filtered MUA (1.2–2.4 Hz for BBN and LED; 1–5 Hz for saccades).

During BBN stimulation, we observed a significant non-random distribution of mean phases in five of the eight recorded brain areas (A1: p = 3 × 10⁻¹^7^; Belt: p = 1 × 10⁻^6^; MGBv: p = 1 × 10⁻⁶; MGB: p = 1 × 10⁻^9^; and mPul: p = 1 × 10⁻⁶). In these regions, which notably are main nodes of the auditory driving pathway, the mean phase corresponded to a high-excitability state, as indicated by the grand average translaminar/transnuclei MUA reaching its positive peak - reflecting elevated neuronal firing - at the time of the BBN bursts (Fig. 9C, blue traces). Moreover, the mean MUA phase distributions across these 5 areas did not differ significantly (Fisher’s test for equality of circular means, p > 0.05), suggesting that excitability was aligned across the auditory thalamocortical network. Such phase alignment likely maximizes the probability of successful signal transmission between auditory-related thalamic and cortical sites by ensuring that interconnected regions reach peak excitability within the same temporal window, thereby facilitating efficient information flow. In contrast, LGN, Pul, and motor cortex, which are part of the auditory modulatory pathway, exhibited a random distribution of mean phases, suggesting that BBN related delta oscillations in these areas did not consistently align to a common excitability phase across trial blocks.

During LED stimulation, pooled mean delta-band MUA phases were significantly clustered in A1 (p = 3 × 10⁻⁶), Belt (p = 2 × 10⁻⁴), MGB (p = 0.01) and mPul (p = 0.01), regions associated with the visual modulatory pathway. Like the pattern observed during BBN stimulation, this clustering occurred around the high-excitability phase, as indicated by the grand average translaminar/transnuclei MUA from these five areas approaching its positive peak at the time of the LED flashes (Fig. 9C, yellow traces). Consistent with coordinated entrainment to the LED stream across these areas, the mean MUA phase distributions did not differ significantly (Fisher’s test for equality of circular means, p > 0.05). Regarding the visual driving pathway, MUA signals in the LGN (p = 5 × 10⁻^4^) and Pul (p = 1 × 10⁻^4^) also exhibited significant phase clustering (Rayleigh’s uniformity test), and were not significantly different neither from each other or the four areas listed above in the visual modulatory pathway (Fisher’s test for equality of circular means, p > 0.05). Mean delta MUA phases from recording sites in MGBv and motor cortex did not show a non-random phase distribution (Rayleigh’s uniformity tests, p > 0.05).

During resting-state saccade-related activity, pooled mean delta/theta MUA phases were significantly clustered in A1 (p = 0.02), LGN (p = 0.002) and mPul (p = 0.004). The angular means of A1 and LGN were nearly identical (3.07 and 2.92 rad, respectively; Fisher’s test for equality of circular means, p = 0.7), whereas the mean phase in mPul was significantly different (–1.33 rad; Fisher’s test, p < 0.0001). These results indicate that saccades entrain delta/theta oscillations in A1 and LGN to a low-excitability phase, as reflected by the grand average translaminar/transnuclear MUA reaching its negative peak at saccade onset (Fig. 9C, pink traces). In contrast, delta/theta oscillations in mPul are entrained to a slightly more excitable phase, with MUA just beginning to approach its positive peak, suggesting a phase offset relative to A1 and LGN. Mean delta/theta MUA phases from the remaining five regions did not exhibit significant clustering (Rayleigh’s tests, p > 0.05).

## DISCUSSION

### Functional differentiation of driving and modulatory circuits

The present study provides direct physiological evidence for a functional distinction between driving and modulatory circuits across cortical and subcortical sensory structures in the non-human primate brain. By jointly quantifying suprathreshold (evoked MUA) and subthreshold (narrowband ITC) activity, we demonstrate that auditory, visual, and motor sampling events can elicit two separable classes of neural responses within an area. Preferred sensory stimuli produce broadband phase alignment, reflected in the uniform area under the curves (AUCs) across adjacent frequency bands (Fig. 6), and are accompanied by significantly increased multiunit firing (Fig. 2). Together, these features are consistent with driving inputs that convey sensory content through strong suprathreshold activation. In contrast, non-preferred sensory and internally generated motor events produced narrowband, frequency-specific ITC modulation - typified by a dominant AUC peak at the rate of stimulation or event occurrence (Figs. 6, 8) - without corresponding increases in firing (Fig. 5). This pattern is consistent with modulatory inputs that regulate ensemble excitability rather than transmit sensory content. These distinctions parallel the conceptual framework of Sherman and Guillery (1998, 2002), who proposed that driving pathways transmit the main information content while modulatory pathways control their efficacy. Our results extend this framework by identifying the spectrotemporal correlates of these circuit types across cortical and thalamic levels in primates. Furthermore, by examining these eight brain structures, we delineate which of these regions are part of a driving, cross-modal- or motor- modulatory circuits (Fig. 5). Additionally, our results reveal that multi-modal events induce coordinated phase alignment of neuronal excitability across cortical and subcortical structures, indicating that rhythmic sensory and motor signals dynamically synchronize network excitability (Fig. 9), which may facilitate efficient cross-modal information flow (e.g. Lakatos et al. 2007).

### Subthreshold modulation as a widespread neural mechanism

A striking feature of the data is the apparent ubiquity of subthreshold modulation across structures and event types. Even in regions not classically associated with the eliciting modality, consistent phase-locked changes in oscillatory activity were observed (see Figs. 5 & 7). This suggests that cross-modal and motor-related signals exert widespread modulatory control, dynamically shaping the excitability of sensory networks. Extensive work in humans and non-human primates has established cross-modal modulation of sensory cortices (reviewed in (Choi, Demir, Oh, & Lee, 2023; Ghazanfar & Schroeder, 2006; Kayser & Logothetis, 2007; Peter Lakatos et al., 2009; Mégevand et al., 2020)) and motor cortex (Chen, Penhune, & Zatorre, 2008; Popescu, Otsuka, & Ioannides, 2004)). Previous primate studies have reported suprathreshold audiovisual integration in the pulvinar (Vittek et al., 2025; Vittek, Juan, Nowak, Girard, & Cappe, 2023), auditory, visual, and somatosensory responses in the superior colliculus (Wallace & Stein, 1994) and human imaging work has identified enhanced LGN and MGB responses during audiovisual co-stimulation (Noesselt et al., 2010). More recently, visual modulation of auditory-related activity was observed in the macaque inferior colliculus (Schmehl, Herche, & Groh, 2025). Subthreshold modulation via phase reset of the local field potential has been posited as a mechanism to mediate such multisensory effects but has generally been studied in the cortex (e.g. Lakatos et al. 2007; 2008; 2009). Our findings extend the subcortical multisensory literature by demonstrating that oscillatory phase alignment is a prominent feature of multisensory processing in subcortical nuclei.

It is well documented in primates that self-generated motor behaviors - such as eye movements, locomotion, and vocalization - modulate neural activity across multiple brain regions, including visual cortex (reviewed in (Gegenfurtner, 2016), auditory cortex (Eliades & Wang, 2003; Leszczynski et al., 2023; O’Connell et al., 2020), somatosensory cortex (Confais, Kim, Tomatsu, Takei, & Seki, 2017; Leszczynski et al., 2023), motor cortex (Cheung, Hamiton, Johnson, & Chang, 2016), LGN (Ramcharan, Gnadt, & Sherman, 2001; Sylvester, Haynes, & Rees, 2005), pulvinar nuclei (Robinson, McClurkin, & Kertzman, 1990; L. Schneider, Dominguez-Vargas, Gibson, Wilke, & Kagan, 2023) and inferior colliculus (Groh, Trause, Underhill, Clark, & Inati, 2001). Consistent with our findings, these studies generally report a suppression of population-level activity associated with self-produced actions (Brooks & Cullen, 2019; Lohse, Zimmer-Harwood, Dahmen, & King, 2022). Our study complements these prior reports by demonstrating that motor-related modulation of sensory activity arises through an excitability reset of low-frequency oscillations. Furthermore, to our knowledge, this is the first evidence that eye movements modulate neuronal activity within the MGB nuclei in primates.

These modulatory effects were most prominent in low-frequency bands, consistent with entrainment to the rates of stimulus presentation (1.6, 1.8, and 6 Hz) and to the saccadic rhythm (∼3 Hz). While a previous study in primate auditory cortex demonstrated entrainment to preferred modality stimuli outside the delta range (Lakatos et al., 2013), our findings extend this work by showing that non-preferred modality stimuli can also entrain neural activity beyond the delta range (i.e. in the theta band). Additionally, we found no significant ITC peaks at frequencies lower than the stimulation rate, indicating that phase resetting does not occur at subharmonics of the stimulation frequency.

### Spectral signatures of driving and modulatory influences

The spectral characteristics of evoked and modulatory responses further clarify their mechanistic distinctions. Consistent with the framework described above, evoked responses exhibited broadband ITC across adjacent frequency bands (Min et al., 2007; Palva, Palva, & Kaila, 2005; Shah et al., 2004). This broadband synchronization likely arises from the synchronous activation of diverse neuronal populations driven by strong, convergent synaptic input. In contrast, previous studies in humans and non-human primates have shown that cross-modal responses that do not elicit suprathreshold activity, instead, elicit modulatory-type responses, which are characterized by frequency-specific phase locking rather than broadband synchronization (Lakatos et al., 2007; Kayser et al., 2008; (Mercier et al., 2013; Thorne, De Vos, Viola, & Debener, 2011). Our findings confirm and extend these reports by demonstrating that modulatory responses in both sensory cortices and subcortical structures, produced by non-preferred sensory and motor events, exhibit phase locking that is tightly aligned with the rate of stimulation for externally driven events (BBN and LED), but extends across neighboring low-frequency bands for internally generated events such as saccades. This pattern suggests that externally driven inputs impose precise temporal structure, whereas internally generated signals engage broader low-frequency coordination. This oscillatory entrainment appears to be mediated by modulatory inputs that shape membrane potential fluctuations without engaging extensive neuronal firing. Such narrowband synchronization may arise through feedback projections from higher-order cortical or thalamic regions, consistent with the known physiology of matrix-type thalamocortical cells and cortico-thalamic modulatory loops (Jones, 2001; Sherman and Guillery, 2011). This aligns with the view that neural communication depends on rhythmic coordination rather than sustained firing ((Engel, Fries, & Singer, 2001; Fries, 2005). Within this framework, driving and modulatory circuits act synergistically: driving inputs convey the content of communication, while modulatory inputs regulate its temporal precision.

### Motor - sensory modulation and predictive timing

Internally generated saccades produced clear modulatory signatures: delta/theta ITC peaks without associated suprathreshold responses. Previous studies in primates have shown that eye movements result in changes in oscillatory activity in sensory cortical areas and hippocampus (Benedetto, Morrone, & Tomassini, 2020; Leszczynski et al., 2025; O’Connell et al., 2020). We add to the literature by showing that eye movements also modulate oscillatory activity in MGB and Pulvinar nuclei. These results suggest that motor-related inputs act primarily through modulatory mechanisms that pre-tune sensory circuits for impending changes in input. The phase resetting induced by saccades may serve to align cortical and thalamic excitability states with expected sensory consequences of movement - a form of predictive timing that minimizes sensory interference during self-generated actions ((Flinker et al., 2010; O’Connell et al., 2020; D. M. Schneider, Nelson, & Mooney, 2014). Additionally, the rebound in excitability following motor inputs is positioned to indirectly enhance sensory processing (Barczak et al., 2019). This interpretation resonates with models of active sensing, in which motor acts synchronize sensory sampling with behaviorally relevant environmental rhythms (Helfrich, 2018; Melloni, Schwiedrzik, Rodriguez, & Singer, 2009; Charles E Schroeder, Wilson, Radman, Scharfman, & Lakatos, 2010). Interestingly, phase alignment across LGN, A1 and mPul following saccades was not perfectly synchronous, with mPul exhibiting a more excitable phase at saccade onset. This pattern suggests that distinct network nodes may experience systematic phase lags consistent with hierarchical processing delays. Supporting this idea, a previous macaque study showed that microstimulation of mPul prior to visual target onset alters saccadic reaction times (Dominguez-Vargas, Schneider, Wilke, & Kagan, 2017), indicating a causal role in shaping saccade timing and visuomotor integration rather than merely reactive processing. Such controlled asynchrony could allow the system to sequentially gate information flow, ensuring that sensory processing unfolds in a temporally ordered manner following each motor event.

## Conclusion

In summary, our findings delineate the physiological organization of driving and modulatory circuits across cortical and subcortical sensory hierarchies. Our results support an integrative model in which driving and modulatory circuits cooperate through oscillatory phase dynamics to regulate sensory processing. Driving inputs convey content via synchronous firing, while modulatory circuits define the temporal windows during which such transmission is effective. Through phase alignment, modulatory pathways can transiently couple regions into coherent processing assemblies, while phase divergence may limit or gate information transfer. This dynamic interplay allows the brain to flexibly route sensory information according to behavioral demands and contextual predictions. By combining measures of suprathreshold and subthreshold activity, our results extend classical anatomical distinctions and link these circuit types to distinct electrophysiological signatures that can be tracked across cortical and thalamic hierarchies. Finally, extending these analyses to behaviorally engaged states could determine how attention or expectation modulate the balance between driving and modulatory activity. Such experiments would provide crucial tests of whether phase coordination serves as an active mechanism for sensory gating and predictive control within the context of driving and modulatory circuit interactions.

## Acknowledgements

Funding sources: R01DC012947 and R01DC019979

## REFERENCES

Barczak, A., Haegens, S., Ross, D. A., McGinnis, T., Lakatos, P., & Schroeder, C. E. (2019). Dynamic Modulation of Cortical Excitability during Visual Active Sensing. Cell reports, 27(12), 3447–3459.e3. doi:10.1016/j.celrep.2019.05.072

Barczak, A., O’Connell, M. N., McGinnis, T., Ross, D., Mowery, T., Falchier, A., & Lakatos, P. (2018). Top-down, contextual entrainment of neuronal oscillations in the auditory thalamocortical circuit. Proceedings of the National Academy of Sciences of the United States of America, 115(32), E7605–E7614. doi:10.1073/pnas.1714684115

Barczak, A., O’Connell, M. N., & Schroeder, C. E. (2023). Thalamic contributions to multisensory convergence and processing. In W. M. Usrey & S. M. Sherman (eds.), The cerebral cortex and thalamus (pp. 305–316). Oxford University PressNew York. doi:10.1093/med/9780197676158.003.0030

Benedetto, A., Morrone, M. C., & Tomassini, A. (2020). The common rhythm of action and perception. Journal of Cognitive Neuroscience, 32(2), 187–200. doi:10.1162/jocn_a_01436

Berg, D. J., Boehnke, S. E., Marino, R. A., Munoz, D. P., & Itti, L. (2009). Free viewing of dynamic stimuli by humans and monkeys. Journal of Vision, 9(5), 19.1–15. doi:10.1167/9.5.19

Besle, J., Schevon, C. A., Mehta, A. D., Lakatos, P., Goodman, R. R., McKhann, G. M., … Schroeder, C. E. (2011). Tuning of the human neocortex to the temporal dynamics of attended events. The Journal of Neuroscience, 31(9), 3176–3185. doi:10.1523/JNEUROSCI.4518-10.2011

Biagioni, T., Fratello, M., Garnier, E., Lagarde, S., Carron, R., Medina Villalon, S., … Pizzo, F. (2025). Interictal waking and sleep electrophysiological properties of the thalamus in focal epilepsies. Brain Communications, 7(2), fcaf102. doi:10.1093/braincomms/fcaf102

Brooks, J. X., & Cullen, K. E. (2019). Predictive sensing: the role of motor signals in sensory processing. Biological Psychiatry: Cognitive Neuroscience and Neuroimaging, 4(9), 842–850. doi:10.1016/j.bpsc.2019.06.003

Brosch, M., Selezneva, E., & Scheich, H. (2005). Nonauditory events of a behavioral procedure activate auditory cortex of highly trained monkeys. The Journal of Neuroscience, 25(29), 6797–6806. doi:10.1523/JNEUROSCI.1571-05.2005

Cappe, C., Rouiller, E. M., & Barone, P. (2009). Multisensory anatomical pathways. Hearing Research, 258(1-2), 28–36. doi:10.1016/j.heares.2009.04.017

Chen, J. L., Penhune, V. B., & Zatorre, R. J. (2008). Listening to musical rhythms recruits motor regions of the brain. Cerebral Cortex, 18(12), 2844–2854. doi:10.1093/cercor/bhn042

Cheung, C., Hamiton, L. S., Johnson, K., & Chang, E. F. (2016). The auditory representation of speech sounds in human motor cortex. eLife, 5. doi:10.7554/eLife.12577

Choi, I., Demir, I., Oh, S., & Lee, S.-H. (2023). Multisensory integration in the mammalian brain: diversity and flexibility in health and disease. Philosophical Transactions of the Royal Society of London. Series B, Biological Sciences, 378(1886), 20220338. doi:10.1098/rstb.2022.0338

Confais, J., Kim, G., Tomatsu, S., Takei, T., & Seki, K. (2017). Nerve-Specific Input Modulation to Spinal Neurons during a Motor Task in the Monkey. The Journal of Neuroscience, 37(10), 2612–2626. doi:10.1523/JNEUROSCI.2561-16.2017

Dominguez-Vargas, A.-U., Schneider, L., Wilke, M., & Kagan, I. (2017). Electrical Microstimulation of the Pulvinar Biases Saccade Choices and Reaction Times in a Time-Dependent Manner. The Journal of Neuroscience, 37(8), 2234–2257. doi:10.1523/JNEUROSCI.1984-16.2016

Eliades, S. J., & Wang, X. (2003). Sensory-motor interaction in the primate auditory cortex during self-initiated vocalizations. Journal of Neurophysiology, 89(4), 2194–2207. doi:10.1152/jn.00627.2002

Engel, A. K., Fries, P., & Singer, W. (2001). Dynamic predictions: oscillations and synchrony in top-down processing. Nature Reviews. Neuroscience, 2(10), 704–716. doi:10.1038/35094565

Flinker, A., Chang, E. F., Kirsch, H. E., Barbaro, N. M., Crone, N. E., & Knight, R. T. (2010). Single-trial speech suppression of auditory cortex activity in humans. The Journal of Neuroscience, 30(49), 16643–16650. doi:10.1523/JNEUROSCI.1809-10.2010

Freeman, J. A., & Nicholson, C. (1975). Experimental optimization of current source-density technique for anuran cerebellum. Journal of Neurophysiology, 38(2), 369–382. doi:10.1152/jn.1975.38.2.369

Fries, P. (2005). A mechanism for cognitive dynamics: neuronal communication through neuronal coherence. Trends in Cognitive Sciences, 9(10), 474–480. doi:10.1016/j.tics.2005.08.011

Fu, K.-M. G., Shah, A. S., O’Connell, M. N., McGinnis, T., Eckholdt, H., Lakatos, P., … Schroeder, C. E. (2004). Timing and laminar profile of eye-position effects on auditory responses in primate auditory cortex. Journal of Neurophysiology, 92(6), 3522–3531. doi:10.1152/jn.01228.2003

Gegenfurtner, K. R. (2016). The interaction between vision and eye movements †. Perception, 45(12), 1333–1357. doi:10.1177/0301006616657097

Ghazanfar, A. A., & Schroeder, C. E. (2006). Is neocortex essentially multisensory? Trends in Cognitive Sciences, 10(6), 278–285. doi:10.1016/j.tics.2006.04.008

Gilbert, C. D., & Li, W. (2013). Top-down influences on visual processing. Nature Reviews. Neuroscience, 14(5), 350–363. doi:10.1038/nrn3476

Groh, J. M., Trause, A. S., Underhill, A. M., Clark, K. R., & Inati, S. (2001). Eye position influences auditory responses in primate inferior colliculus. Neuron, 29(2), 509–518. doi:10.1016/s0896-6273(01)00222-7

Haegens, S., & Zion Golumbic, E. (2018). Rhythmic facilitation of sensory processing: A critical review. Neuroscience and Biobehavioral Reviews, 86, 150–165. doi:10.1016/j.neubiorev.2017.12.002

Helfrich, R. F. (2018). The rhythmic nature of visual perception. Journal of Neurophysiology, 119(4), 1251–1253. doi:10.1152/jn.00810.2017

Helfrich, R. F., & Knight, R. T. (2016). Oscillatory dynamics of prefrontal cognitive control. Trends in Cognitive Sciences, 20(12), 916–930. doi:10.1016/j.tics.2016.09.007

Hoffman, K. L., Dragan, M. C., Leonard, T. K., Micheli, C., Montefusco-Siegmund, R., & Valiante, T. A. (2013). Saccades during visual exploration align hippocampal 3-8 Hz rhythms in human and non-human primates. Frontiers in Systems Neuroscience, 7, 43. doi:10.3389/fnsys.2013.00043

Hubel, D. H., & Wiesel, T. N. (1962). Receptive fields, binocular interaction and functional architecture in the cat’s visual cortex. The Journal of Physiology, 160, 106–154. doi:10.1113/jphysiol.1962.sp006837

Ito, J., Maldonado, P., & Grün, S. (2013). Cross-frequency interaction of the eye-movement related LFP signals in V1 of freely viewing monkeys. Frontiers in Systems Neuroscience, 7, 1. doi:10.3389/fnsys.2013.00001

Ito, J., Maldonado, P., Singer, W., & Grün, S. (2011). Saccade-related modulations of neuronal excitability support synchrony of visually elicited spikes. Cerebral Cortex, 21(11), 2482–2497. doi:10.1093/cercor/bhr020

Jones, E. G. (1998). Viewpoint: the core and matrix of thalamic organization. Neuroscience, 85(2), 331–345. doi:10.1016/s0306-4522(97)00581-2

Jones, E. G. (2001). The thalamic matrix and thalamocortical synchrony. Trends in Neurosciences, 24(10), 595–601. doi:10.1016/s0166-2236(00)01922-6

Kajikawa, Y., & Schroeder, C. E. (2011). How local is the local field potential? Neuron, 72(5), 847–858. doi:10.1016/j.neuron.2011.09.029

Kajikawa, Y., & Schroeder, C. E. (2015). Generation of field potentials and modulation of their dynamics through volume integration of cortical activity. Journal of Neurophysiology, 113(1), 339–351. doi:10.1152/jn.00914.2013

Kayser, C., & Logothetis, N. K. (2007). Do early sensory cortices integrate cross-modal information? Brain Structure & Function, 212(2), 121–132. doi:10.1007/s00429-007-0154-0

Kayser, C., Petkov, C. I., & Logothetis, N. K. (2008). Visual modulation of neurons in auditory cortex. Cerebral Cortex, 18(7), 1560–1574. doi:10.1093/cercor/bhm187

Lakatos, P, Chen, C. M., O’Connell, M. N., Mills, A., & Schroeder, C. E. (2007). Neuronal oscillations and multisensory interaction in primary auditory cortex. Neuron, 53(2), 279–292. doi:10.1016/j.neuron.2006.12.011

Lakatos, Peter, Karmos, G., Mehta, A. D., Ulbert, I., & Schroeder, C. E. (2008). Entrainment of neuronal oscillations as a mechanism of attentional selection. Science, 320(5872), 110–113. doi:10.1126/science.1154735

Lakatos, Peter, Musacchia, G., O’Connel, M. N., Falchier, A. Y., Javitt, D. C., & Schroeder, C. E. (2013). The spectrotemporal filter mechanism of auditory selective attention. Neuron, 77(4), 750–761. doi:10.1016/j.neuron.2012.11.034

Lakatos, Peter, O’Connell, M. N., Barczak, A., McGinnis, T., Neymotin, S., Schroeder, C. E., … Javitt, D. C. (2020). The thalamocortical circuit of auditory mismatch negativity. Biological Psychiatry, 87(8), 770–780. doi:10.1016/j.biopsych.2019.10.029

Lakatos, Peter, O’Connell, M. N., Barczak, A., Mills, A., Javitt, D. C., & Schroeder, C. E. (2009). The leading sense: supramodal control of neurophysiological context by attention. Neuron, 64(3), 419–430. doi:10.1016/j.neuron.2009.10.014

Lakatos, Peter, Pincze, Z., Fu, K.-M. G., Javitt, D. C., Karmos, G., & Schroeder, C. E. (2005). Timing of pure tone and noise-evoked responses in macaque auditory cortex. Neuroreport, 16(9), 933–937. doi:10.1097/00001756-200506210-00011

Lakatos, Peter, Shah, A. S., Knuth, K. H., Ulbert, I., Karmos, G., & Schroeder, C. E. (2005). An oscillatory hierarchy controlling neuronal excitability and stimulus processing in the auditory cortex. Journal of Neurophysiology, 94(3), 1904–1911. doi:10.1152/jn.00263.2005

Lee, D., & Malpeli, J. G. (1998). Effects of saccades on the activity of neurons in the cat lateral geniculate nucleus. Journal of Neurophysiology, 79(2), 922–936. doi:10.1152/jn.1998.79.2.922

Legatt, A. D., Arezzo, J., & Vaughan, H. G. (1980). Averaged multiple unit activity as an estimate of phasic changes in local neuronal activity: effects of volume-conducted potentials. Journal of Neuroscience Methods, 2(2), 203–217. doi:10.1016/0165-0270(80)90061-8

Leszczynski, M., Bickel, S., Nentwich, M., Russ, B. E., Parra, L., Lakatos, P., … Schroeder, C. E. (2023). Saccadic modulation of neural excitability in auditory areas of the neocortex. Current Biology, 33(7), 1185–1195.e6. doi:10.1016/j.cub.2023.02.018

Leszczynski, M., Espinal, E., Smith, E., Schevon, C., Sheth, S., & Schroeder, C. E. (2025). Eye movements organize excitability state, information coding and network connectivity in the human hippocampus. Progress in Neurobiology, 252, 102812. doi:10.1016/j.pneurobio.2025.102812

Lohse, M., Zimmer-Harwood, P., Dahmen, J. C., & King, A. J. (2022). Integration of somatosensory and motor-related information in the auditory system. Frontiers in Neuroscience, 16, 1010211. doi:10.3389/fnins.2022.1010211

Mackey, C. A., O’Connell, M. N., Hackett, T. A., Schroeder, C. E., & Kajikawa, Y. (2024). Laminar organization of visual responses in core and parabelt auditory cortex. Cerebral Cortex, 34(9). doi:10.1093/cercor/bhae373

Maldonado, P., Babul, C., Singer, W., Rodriguez, E., Berger, D., & Grün, S. (2008). Synchronization of neuronal responses in primary visual cortex of monkeys viewing natural images. Journal of Neurophysiology, 100(3), 1523–1532. doi:10.1152/jn.00076.2008

Mégevand, P., Mercier, M. R., Groppe, D. M., Zion Golumbic, E., Mesgarani, N., Beauchamp, M. S., … Mehta, A. D. (2020). Crossmodal phase reset and evoked responses provide complementary mechanisms for the influence of visual speech in auditory cortex. The Journal of Neuroscience, 40(44), 8530–8542. doi:10.1523/JNEUROSCI.0555-20.2020

Melloni, L., Schwiedrzik, C. M., Rodriguez, E., & Singer, W. (2009). (Micro)Saccades, corollary activity and cortical oscillations. Trends in Cognitive Sciences, 13(6), 239–245. doi:10.1016/j.tics.2009.03.007

Mercier, M. R., Foxe, J. J., Fiebelkorn, I. C., Butler, J. S., Schwartz, T. H., & Molholm, S. (2013). Auditory-driven phase reset in visual cortex: human electrocorticography reveals mechanisms of early multisensory integration. Neuroimage, 79, 19–29. doi:10.1016/j.neuroimage.2013.04.060

Merzenich, M. M., & Brugge, J. F. (1973). Representation of the cochlear partition of the superior temporal plane of the macaque monkey. Brain Research, 50(2), 275–296. doi:10.1016/0006-8993(73)90731-2

Min, B.-K., Busch, N. A., Debener, S., Kranczioch, C., Hanslmayr, S., Engel, A. K., & Herrmann, C. S. (2007). The best of both worlds: phase-reset of human EEG alpha activity and additive power contribute to ERP generation. International Journal of Psychophysiology, 65(1), 58–68. doi:10.1016/j.ijpsycho.2007.03.002

Mitzdorf, U. (1985). Current source-density method and application in cat cerebral cortex: investigation of evoked potentials and EEG phenomena. Physiological Reviews, 65(1), 37–100. doi:10.1152/physrev.1985.65.1.37

Mountcastle, V. B. (1957). Modality and topographic properties of single neurons of cat’s somatic sensory cortex. Journal of Neurophysiology, 20(4), 408–434. doi:10.1152/jn.1957.20.4.408

Naue, N., Rach, S., Strüber, D., Huster, R. J., Zaehle, T., Körner, U., & Herrmann, C. S. (2011). Auditory event-related response in visual cortex modulates subsequent visual responses in humans. The Journal of Neuroscience, 31(21), 7729–7736. doi:10.1523/JNEUROSCI.1076-11.2011

Nicholson, C., & Freeman, J. A. (1975). Theory of current source-density analysis and determination of conductivity tensor for anuran cerebellum. Journal of Neurophysiology, 38(2), 356–368. doi:10.1152/jn.1975.38.2.356

Noesselt, T., Tyll, S., Boehler, C. N., Budinger, E., Heinze, H.-J., & Driver, J. (2010). Sound-induced enhancement of low-intensity vision: multisensory influences on human sensory-specific cortices and thalamic bodies relate to perceptual enhancement of visual detection sensitivity. The Journal of Neuroscience, 30(41), 13609–13623. doi:10.1523/JNEUROSCI.4524-09.2010

O’Connell, M. N., Barczak, A., McGinnis, T., Mackin, K., Mowery, T., Schroeder, C. E., & Lakatos, P. (2020). The role of motor and environmental visual rhythms in structuring auditory cortical excitability. iScience, 23(8), 101374. doi:10.1016/j.isci.2020.101374

O’Connell, M. N., Falchier, A., McGinnis, T., Schroeder, C. E., & Lakatos, P. (2011). Dual mechanism of neuronal ensemble inhibition in primary auditory cortex. Neuron, 69(4), 805–817. doi:10.1016/j.neuron.2011.01.012

Obleser, J., & Kayser, C. (2019). Neural entrainment and attentional selection in the listening brain. Trends in Cognitive Sciences, 23(11), 913–926. doi:10.1016/j.tics.2019.08.004

Palva, J. M., Palva, S., & Kaila, K. (2005). Phase synchrony among neuronal oscillations in the human cortex. The Journal of Neuroscience, 25(15), 3962–3972. doi:10.1523/JNEUROSCI.4250-04.2005

Plass, J., Ahn, E., Towle, V. L., Stacey, W. C., Wasade, V. S., Tao, J., … Brang, D. (2019). Joint encoding of auditory timing and location in visual cortex. Journal of Cognitive Neuroscience, 31(7), 1002–1017. doi:10.1162/jocn_a_01399

Popescu, M., Otsuka, A., & Ioannides, A. A. (2004). Dynamics of brain activity in motor and frontal cortical areas during music listening: a magnetoencephalographic study. Neuroimage, 21(4), 1622–1638. doi:10.1016/j.neuroimage.2003.11.002

Ramcharan, E. J., Gnadt, J. W., & Sherman, S. M. (2001). The effects of saccadic eye movements on the activity of geniculate relay neurons in the monkey. Visual Neuroscience, 18(2), 253–258. doi:10.1017/s0952523801182106

Rauschecker, J. P., Tian, B., Pons, T., & Mishkin, M. (1997). Serial and parallel processing in rhesus monkey auditory cortex. The Journal of Comparative Neurology, 382(1), 89–103. doi:10.1002/(SICI)1096-9861(19970526)382:1<89::AID-CNE6>3.0.CO;2-G

Rayner, K. (1998). Eye movements in reading and information processing: 20 years of research. Psychological Bulletin, 124(3), 372–422. doi:10.1037/0033-2909.124.3.372

Reppas, J. B., Usrey, W. M., & Reid, R. C. (2002). Saccadic eye movements modulate visual responses in the lateral geniculate nucleus. Neuron, 35(5), 961–974. doi:10.1016/s0896-6273(02)00823-1

Robinson, D. L., McClurkin, J. W., & Kertzman, C. (1990). Orbital position and eye movement influences on visual responses in the pulvinar nuclei of the behaving macaque. Experimental Brain Research, 82(2), 235–246. doi:10.1007/BF00231243

Romei, V., Gross, J., & Thut, G. (2012). Sounds reset rhythms of visual cortex and corresponding human visual perception. Current Biology, 22(9), 807–813. doi:10.1016/j.cub.2012.03.025

Schmehl, M. N., Herche, J. L., & Groh, J. M. (2025). Visually evoked activity and variable modulation of auditory responses in the macaque inferior colliculus. Journal of Neurophysiology, 133(5), 1456–1467. doi:10.1152/jn.00529.2024

Schneider, D. M., Nelson, A., & Mooney, R. (2014). A synaptic and circuit basis for corollary discharge in the auditory cortex. Nature, 513(7517), 189–194. doi:10.1038/nature13724

Schneider, L., Dominguez-Vargas, A.-U., Gibson, L., Wilke, M., & Kagan, I. (2023). Visual, delay, and oculomotor timing and tuning in macaque dorsal pulvinar during instructed and free choice memory saccades. Cerebral Cortex, 33(21), 10877–10900. doi:10.1093/cercor/bhad333

Schroeder, C E, Lindsley, R. W., Specht, C., Marcovici, A., Smiley, J. F., & Javitt, D. C. (2001). Somatosensory input to auditory association cortex in the macaque monkey. Journal of Neurophysiology, 85(3), 1322–1327. doi:10.1152/jn.2001.85.3.1322

Schroeder, C E, Mehta, A. D., & Givre, S. J. (1998). A spatiotemporal profile of visual system activation revealed by current source density analysis in the awake macaque. Cerebral Cortex, 8(7), 575–592. doi:10.1093/cercor/8.7.575

Schroeder, Charles E, & Lakatos, P. (2009). Low-frequency neuronal oscillations as instruments of sensory selection. Trends in Neurosciences, 32(1), 9–18. doi:10.1016/j.tins.2008.09.012

Schroeder, Charles E, Wilson, D. A., Radman, T., Scharfman, H., & Lakatos, P. (2010). Dynamics of Active Sensing and perceptual selection. Current Opinion in Neurobiology, 20(2), 172–176. doi:10.1016/j.conb.2010.02.010

Shah, A. S., Bressler, S. L., Knuth, K. H., Ding, M., Mehta, A. D., Ulbert, I., & Schroeder, C. E. (2004). Neural dynamics and the fundamental mechanisms of event-related brain potentials. Cerebral Cortex, 14(5), 476–483. doi:10.1093/cercor/bhh009

Sherman, S. M., & Guillery, R. W. (2002). The role of the thalamus in the flow of information to the cortex. Philosophical Transactions of the Royal Society of London. Series B, Biological Sciences, 357(1428), 1695–1708. doi:10.1098/rstb.2002.1161

Sherman, S. M., & Guillery, R. W. (2011). Distinct functions for direct and transthalamic corticocortical connections. Journal of Neurophysiology, 106(3), 1068–1077. doi:10.1152/jn.00429.2011

Steinschneider, M., Reser, D., Schroeder, C. E., & Arezzo, J. C. (1995). Tonotopic organization of responses reflecting stop consonant place of articulation in primary auditory cortex (A1) of the monkey. Brain Research, 674(1), 147–152. doi:10.1016/0006-8993(95)00008-e

Sylvester, R., Haynes, J.-D., & Rees, G. (2005). Saccades differentially modulate human LGN and V1 responses in the presence and absence of visual stimulation. Current Biology, 15(1), 37–41. doi:10.1016/j.cub.2004.12.061

Thorne, J. D., De Vos, M., Viola, F. C., & Debener, S. (2011). Cross-modal phase reset predicts auditory task performance in humans. The Journal of Neuroscience, 31(10), 3853–3861. doi:10.1523/JNEUROSCI.6176-10.2011

Vittek, A.-L., Juan, C., Gaillard, C., Mercier, M., Girard, P., Ben Hamed, S., & Cappe, C. (2025). Frequency coding of multisensory integration in the local field potentials of the medial pulvinar. The European Journal of Neuroscience, 62(5), e70230. doi:10.1111/ejn.70230

Vittek, A.-L., Juan, C., Nowak, L. G., Girard, P., & Cappe, C. (2023). Multisensory integration in neurons of the medial pulvinar of macaque monkey. Cerebral Cortex, 33(8), 4202–4215. doi:10.1093/cercor/bhac337

Wallace, M. T., & Stein, B. E. (1994). Cross-modal synthesis in the midbrain depends on input from cortex. Journal of Neurophysiology, 71(1), 429–432. doi:10.1152/jn.1994.71.1.429

